# Estrogen exacerbates mammary involution through neutrophil dependent and independent mechanism

**DOI:** 10.1101/2020.04.03.023341

**Authors:** Chew Leng Lim, Yu Zuan Or, Zoe Ong, Hwa Hwa Chung, Hirohito Hayashi, Smeeta Shrestha, Shunsuke Chiba, Lin Feng, Valerie CL Lin

**Affiliations:** NTU Institute for Health Technologies, Interdisciplinary Graduate School, Nanyang Technological University, Singapore; School of Biological Sciences, Nanyang Technological University, Singapore; Division of Chemistry and Biological Chemistry, School of Physical and Mathematical Sciences, Nanyang Technological University, Singapore; School of Computer Science and Engineering, Nanyang Technological University, Singapore; School of Basic and Applied Sciences, Dayananda Sagar University, Bangalore, India

## Abstract

There is strong evidence that the pro-inflammatory microenvironment during post-partum mammary involution promotes parity-associated breast cancer. Estrogen exposure during mammary involution drives tumour growth through the activity of neutrophils. However, how estrogen and neutrophils influence mammary involution are unknown. Combined analysis of transcriptomic, protein, and immunohistochemical data in Balb/c mice with and without neutrophil depletion showed that estrogen promotes involution by exacerbating inflammation, cell death and adipocytes repopulation through neutrophil-dependent and neutrophil-independent mechanisms. Remarkably, 88% of estrogen-regulated genes in mammary tissue were mediated through neutrophils, which were recruited through estrogen-induced CXCL2-CXCR2 signalling. While neutrophils mediate estrogen-induced inflammation and adipocytes repopulation, estrogen-induced mammary cell death was mediated by neutrophils-independent upsurges of cathepsins and their lysosomal leakages that are critical for lysosome-mediated cell death. Notably, these multifaceted effects of estrogen are unique to the phase of mammary involution. These findings are important for the development of intervention strategies for parity-associated breast cancer.

## Introduction

There is strong evidence that the mammary microenvironment during the post-partum mammary involution promotes mammary tumour progression. High levels of tissue fibrillar collagen and elevated expression of cyclooxygenase-2 (COX-2) in the mammary gland have been shown to drive tumour growth and lymph angiogenesis [1, 2]. Wound healing-like tissue environment associated with mammary involution is also known to promote tumour development and dissemination [3]. Estrogen has been shown to stimulate the growth of estrogen receptor-negative mammary tumours during mammary involution and estrogen-stimulated neutrophil activity plays a crucial role in fostering the pro-tumoral microenvironment [4]. This suggests that estrogen exposure during post-weaning mammary involution is a risk factor for parity-associated breast cancer. However, the functional roles of estrogen and neutrophils in mammary biology during involution have been little studied to date.

Post-weaning mammary involution is a process for the lactating mammary gland to return to the pre-pregnancy state. The distinctive features of mammary involution include massive cell death of the secretory mammary alveoli, acute inflammation, extracellular matrix remodelling and adipocyte repopulation. Involution is commonly divided into two phases. In mice, the first phase is the reversible phase whereby the reintroduction of the pups within 48h can re-initiate lactation [5, 6]. It is typified by a decrease in milk protein synthesis and increased mammary cell death resulting in the appearance of shed, dying cells within the lumen of the distended alveoli [7, 8]. Inflammation also occurs in the first phase with the infiltration of immune cells and up-regulation of immune response genes [9, 10]. The second, and irreversible phase of mammary involution occurs after 72h of weaning. This phase is morphologically characterized by the collapse of the alveolar structure, the second wave of epithelial cell death, continued inflammation and the repopulation of adipocytes. This is followed by mammary regeneration and tissue remodelling to the pre-pregnancy state.

Neutrophils are the most abundant leukocytes in the innate immune system against invading pathogens. Neutrophils are also activated in response to sterile inflammation, but the outcome is complicated and depends on the context [11]. The importance of neutrophils in tumour development has been increasingly recognized in recent years. Tumour-associated neutrophils have been classified into the anti-tumour neutrophils (N1) and pro-tumour neutrophils (N2) based on the expression of specific markers [12]. Neutrophils are known to accumulate in the peripheral blood of patients with cancer and a high circulating neutrophil-to-lymphocyte ratio is known as a strong biomarker of poor prognosis in various cancers [13]. Based on density gradient centrifugation, circulating neutrophils in mice model can also be classified into the low-density and high-density neutrophils (LDN and HDN) [14, 15]. LDN was associated with immunosuppressive activity promoting tumour growth while HDN was considered cytotoxic to the tumour. Neutrophils were reported to be the first immune cells recruited into mammary tissue during involution [9], although the significance of their presence is not known.

Estrogen is well studied on its mitogenic effect and plays essential roles in mammary ductal development. However, the studies of estrogen influence on mammary involution are scarce. In mice, a histological examination in 1970 reported that estrogen treatment retards mammary involution [16]. However, estrogen was reported to promote the regression of mammary glands in rats by up-regulating a 60K gelatinase which degrades collagen and participate in extracellular matrix remodelling [17]. Similar to the observation in rats, injection of estrogen during the dry off period in dairy cows hastened involution based on the abrupt decline of milk synthesis and secretion [18, 19]. Those early preliminary studies clearly indicate the involvement of estrogen in the modulation of mammary involution, but the pieces of evidence are limited and inconsistent among different models.

In this present study, we investigated the influence of estrogen on the progression of mammary involution in mice. We also evaluated the roles of neutrophils in estrogen regulation of this process in neutrophil depletion experiments. Our data show that estrogen plays a multifaceted role in the regulation of post-weaning mammary involution by enhancing inflammation, cell death, adipocyte occupancy, and tissue remodelling through both neutrophil-dependent and neutrophil-independent mechanisms. Global transcriptomic analysis revealed striking effect of estrogen on neutrophil activities that likely exert profound influence on mammary microenvironment that may have significantly impact on the development of parity-associated breast cancer.

## Result

### Estrogen promotes inflammation, programmed cell death and adipocytes repopulation during post-weaning mammary involution

Acute inflammation, programmed cell death, and adipocytes repopulation are the hallmarks of the acute phase of post-weaning mammary involution. To understand the roles of estrogen in mammary involution, its effect on these hallmarks were evaluated. Ovariectomized (OVX) mice at 24h post-weaning (INV D1) were treated with or without 17β-estradiol benzoate (E2B) for 48h. E2B caused significantly more mammary cell death, as evidenced by the increase of the number of shed cells with hyper-condensed nuclei in the lumen, a characteristic of programmed cell death (Fig. 1, Ai, and 1B, p=0.0131). This effect was associated with an increase in the number of cleaved caspase-3-positive (CC3^+^) mammary cells (Fig. 1, Aii, and 1B, p=0.0283). E2B also significantly increased the repopulation of adipocytes based on the staining of perilipin, a lipid droplet-associated protein [20] (Fig. 1, Aiii, and 1B, p=0.0155). Accordingly, E2B decreased the expression of milk proteins β-casein (*Csn2*) (Fig. 1C, p=0.0187).

**Figure 1.**
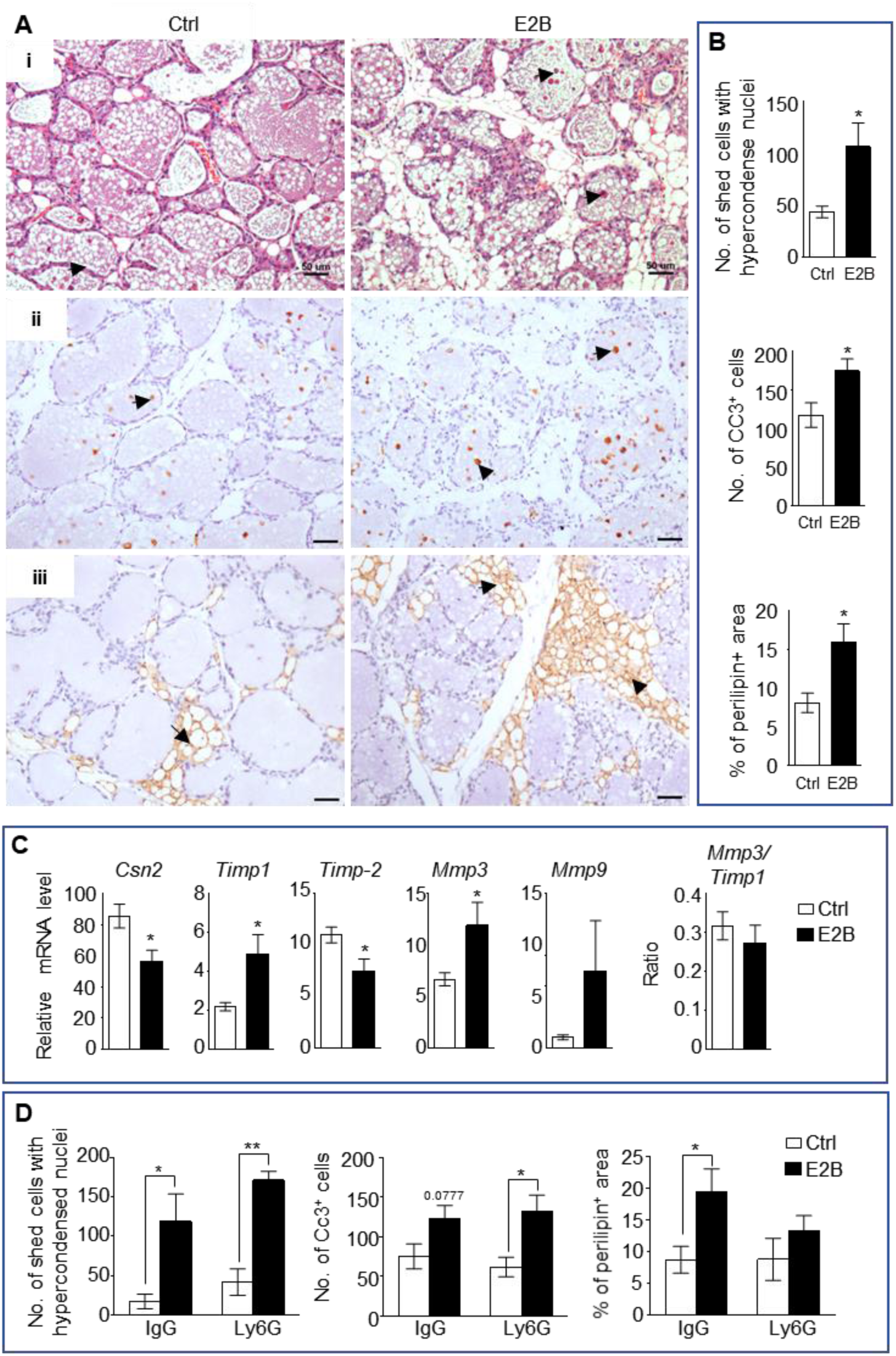
Estrogen accelerates mammary involution. Mice on the day of weaning (involution day 1-INV D1) were treated with vehicle control (Ctrl) or E2B for 48h before mammary tissues were collected for analysis. A, (i) H&E stained mammary tissue sections; shed cells with hyper-condensed nuclei are indicated by arrows. (ii) IHC of cleaved caspase-3 (CC3); arrows indicate CC3^+^ cells. (iii) Perilipin IHC; arrows indicate perilipin^+^ adipocytes. Scale bars: 50µm. B, Quantification of the number of shed cells with hypercondensed nuclei in the lumens from H&E sections (Ctrl n=9, E2B n=8), of number of CC3^+^ cells (Ctrl n=7, E2B n=6), and of percentage of perilipin stained area (Ctrl n=7, E2B n=8). C, Gene expression of *Csn2* and tissue remodelling enzymes *Timp1, Timp2, Mmp3, and Mmp9* relative to *36b4* by qPCR analysis (Ctrl n=7, E2B n=6). D, Neutrophil depletion had no effect on cell shedding and number of CC3^+^ cells, but attenuated estrogen-induced adipocytes repopulation (Ctrl+IgG n=4, E2B+IgG n=4, Ctrl+Ly6G n=3, E2B+Ly6G, n=3). Data represented as mean ± SEM.

We reported previously that E2B markedly induced the expression of inflammatory genes and neutrophil infiltration [4]. We questioned if neutrophils are involved in E2B-induced cell death and adipocytes repopulation. Interestingly, neutrophil depletion had no effect on E2B-induced mammary cell death but diminished E2B-induced adipocytes repopulation (Fig. 1D). The representative histological images of the effect of neutrophil depletion are shown in Suppl. Fig. 2. Taken together, we conclude that estrogen promotes mammary involution, and neutrophils are critical for estrogen-induced inflammation and adipocytes repopulation, but not for mammary cell death.

### Majority of estrogen-regulated genes in involuting mammary tissue is mediated through neutrophils

To elucidate the mechanism of estrogen regulation during mammary involution, RNA-Seq analysis of mammary tissue was conducted. Since estrogen has been shown to induce neutrophil infiltration in mammary tissue, the involvement of neutrophils in estrogen regulation of gene expression was also determined. OVX mice were injected with anti-Ly6G antibody (Ly6G) to deplete neutrophils at 24h post-weaning (INV D1). Mice given isotype control antibody (IgG) were used as controls. Mice were subsequently treated with or without E2B for 24h. Consistent with the previous study [4], E2B induced neutrophil infiltration during mammary involution. E2B also induced an increase of mammary tissue monocytes (CD45+ CD11b+ Ly6C^hi^), which can be considered as mammary macrophages because they have infiltrated into the tissue. Anti-Ly6G treatment reduced neutrophil (CD45+ CD11b+ Gr1^hi^) levels in the blood and mammary tissue by more than 95% (Suppl. Fig. 1, p<0.05). Mammary macrophages were decreased with neutrophil depletion by anti-Ly6G, although this reduction was not statistically significant (Suppl. Fig. 1B, p=0.1417). It is plausible that E2B-induced monocytes infiltration is partly mediated by neutrophils.

RNA-Seq data were analysed using DESeq2 [21] to identify differentially expressed (DE) genes between the different treatments. As shown in the volcano plot (Fig. 2A, top panel), a total of 1987 genes were significantly (padj<0.05) regulated by E2B in IgG groups (Fig. 2B) with very high fold changes. Of these genes, 513 genes were up-regulated, and 148 genes were down-regulated with a fold change ≥ 2 (Fig. 2A, top panel). Gene ontology (GO) analysis showed that estrogen regulates genes involved in diverse biological processes (Fig. 2C). Leukocyte migration, regulation of inflammatory response, and cytokine-mediated signalling are among the top biological processes regulated. E2B also regulated genes in epithelial cell proliferation, cell death, and fat cell differentiation.

**Figure 2.**
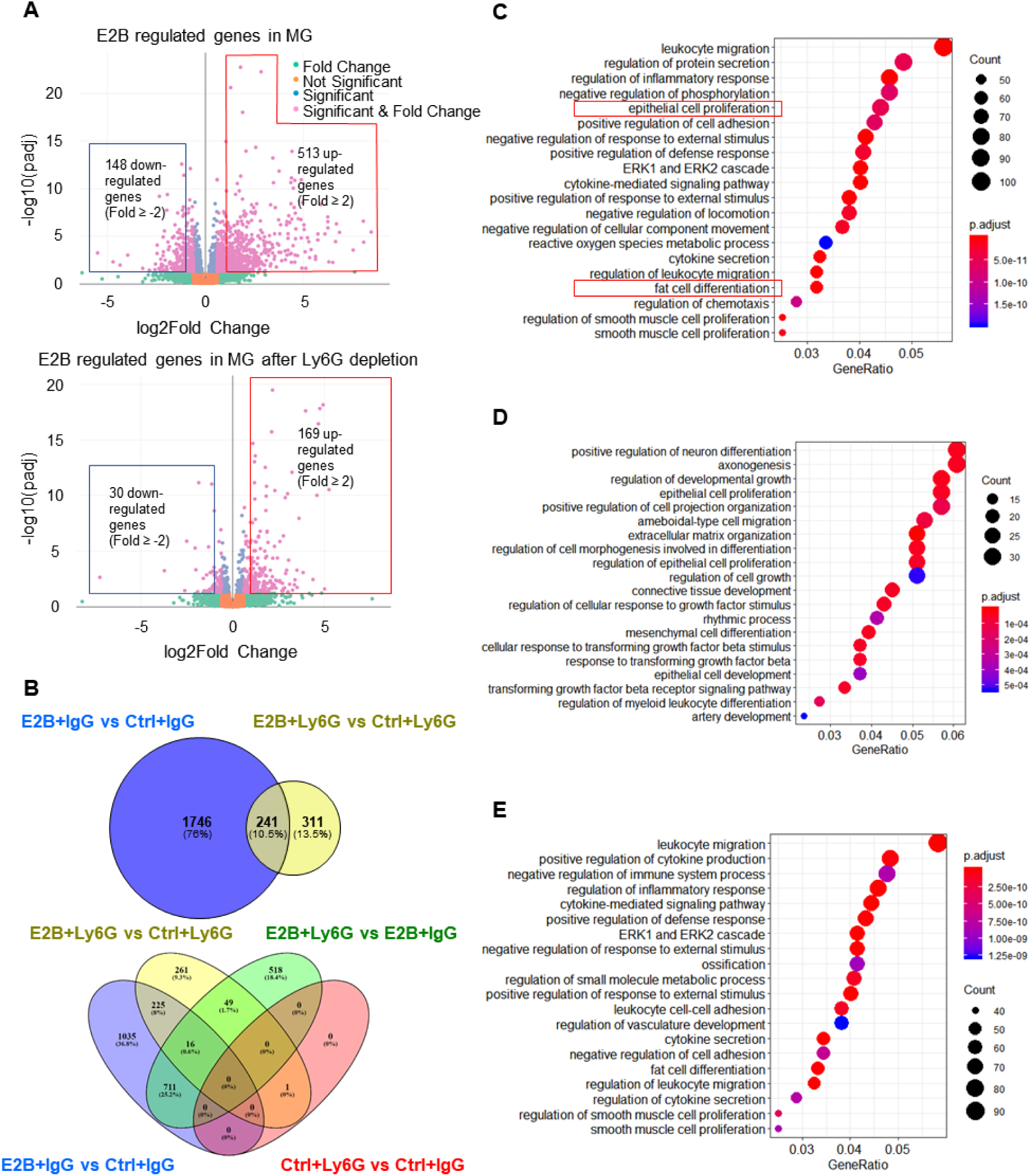
Estrogen regulates a multitude of neutrophil-dependent and –independent biological processes in involuting mammary gland. Mice at INV D1 were treated with anti-Ly6G antibody (Ly6G) or isotype control (IgG). 24h later, they were treated with vehicle control (Ctrl) or E2B for 24h (Ctrl+IgG n=3, Ctrl+Ly6G n=3, E2B+IgG n=3, E2B+Ly6G n=3). RNA-Seq data were processed and analysed with DESeq2 followed by GO over-representation analysis. A, Volcano plot for the differentially expressed E2B regulated genes in mammary gland (MG) from IgG- and Ly6G-treated animals. B, Venn diagram for the differentially expressed genes identified from the DESeq2 analysis of the RNA-Seq data. C, Top 20 Gene Ontology (GO) terms for E2B regulated genes in MG without neutrophil depletion. D, Top 20 GO terms for the E2B regulated genes in MG after neutrophil depletion. E, Top 20 GO terms for E2B regulated genes are lost as a result of neutrophil depletion.

Remarkably, neutrophil depletion eliminated 88% of estrogen-regulated genes (Fig. 2B, top panel), i.e., 1746 out of 1987 E2B-regulated genes in mammary tissue were regulated in neutrophils directly, or indirectly through regulating the activities of neutrophils. Correspondingly, 12% (241 genes) of E2B-regulated genes in mammary tissue are independent of neutrophils. It is also noteworthy that there is only one differentially regulated gene (padj<0.05) between IgG and Ly6G groups in the absence of estrogen (Fig. 2B, bottom panel). This could be due to the fact that the number of mammary neutrophils present in these Ctrl samples were too low (less than 1% of the total live cells in FACS analysis) to account for any differences in gene expression. Consistent with the function of neutrophils, the GO terms including leukocyte migration, inflammatory response, cytokines and chemokines related pathways, and fat cell differentiation that were regulated by E2B were all eliminated in neutrophil depleted samples (Fig. 2D). On the other hand, GO terms such as epithelial cell proliferation and developmental growth persisted in neutrophil-depleted samples suggesting that their stimulation by E2B are independent of neutrophils (Fig. 2C, 2D).

Taken together, global gene expression analysis of mammary tissue demonstrates that estrogen regulates genes in a plethora of biological processes during post-weaning mammary involution, and 88% of estrogen-regulated genes are mediated through neutrophils. Therefore, the E2B-induced pro-inflammatory microenvironment during mammary involution is primarily due to its regulation of neutrophil activities, which play a significant role in post-weaning mammary involution. In the subsequent studies, we investigated the mechanism of estrogen-induced neutrophil infiltration, mammary cell death, and adipocytes repopulation during post-weaning mammary involution.

### Estrogen-induced *Cxcr2* signalling in neutrophils plays a key role in mammary neutrophil infiltration

Based on the GO over-representation analysis, 63 E2B-regulated genes with fold change ≥ 3 were associated with leukocyte migration and inflammation (Fig. 3A). All 63 genes were no longer E2B-regulated after neutrophil depletion (in Ly6G group). This suggests that the regulation of inflammation by estrogen is primarily exerted through neutrophils. Consistent with the previous report [4], S100 calcium-binding protein A8 (*S100a8*) and A9 (*S100a9*) were all induced by E2B in isolated mammary neutrophils using magnetic beads (Dynabeads®) coupled to anti-Ly6G antibody (Fig. 3B, *S100a8*, p=0.0079; *S100a9*, p=0.0035). C-X-C motif chemokine receptor 2 (*Cxcr2*), the receptor for C-X-C motif chemokine ligand 1 and 2 (*Cxcl1* and *Cxcl2*) which was also previously demonstrated to be induced by estrogen in mammary neutrophils [4], was also significantly up-regulated (Fig. 3B, p=0.0139). Aconitate decarboxylase 1 (*Acod1*), triggering receptor expressed on myeloid cells 1 (*Trem1*), triggering receptor expressed on myeloid cells 3 (*Trem3*), and interleukin 1 beta (*Il1b*) that displayed high fold induction were also validated with qPCR in isolated neutrophils (Fig. 3B, *Acod1*, p=0.0845; *Trem1*, p=0.0252; *Trem3*, p=0.0181; *Il1b*, p=0.0527). The results confirm that the up-regulation of inflammatory genes was a result of estrogen induction in mammary neutrophils. Another interesting estrogen-regulated gene is myocardial infraction associated transcript 2 (*Mirt2*; Fig. 3B, p=0.008), which was found to be one of the highest E2B-regulated gene in the RNA-Seq data. *Mirt2* is a lipopolysaccharides (LPS)-induced long non-coding RNA (lncRNA) in macrophages and was reported to be a negative regulator of LPS-induced inflammation both *in vivo* and *in vitro* [22]. Thus, *Mirt2* regulation by E2B in neutrophils indicates its involvement in the moderation of the inflammatory response during mammary involution.

**Figure 3.**
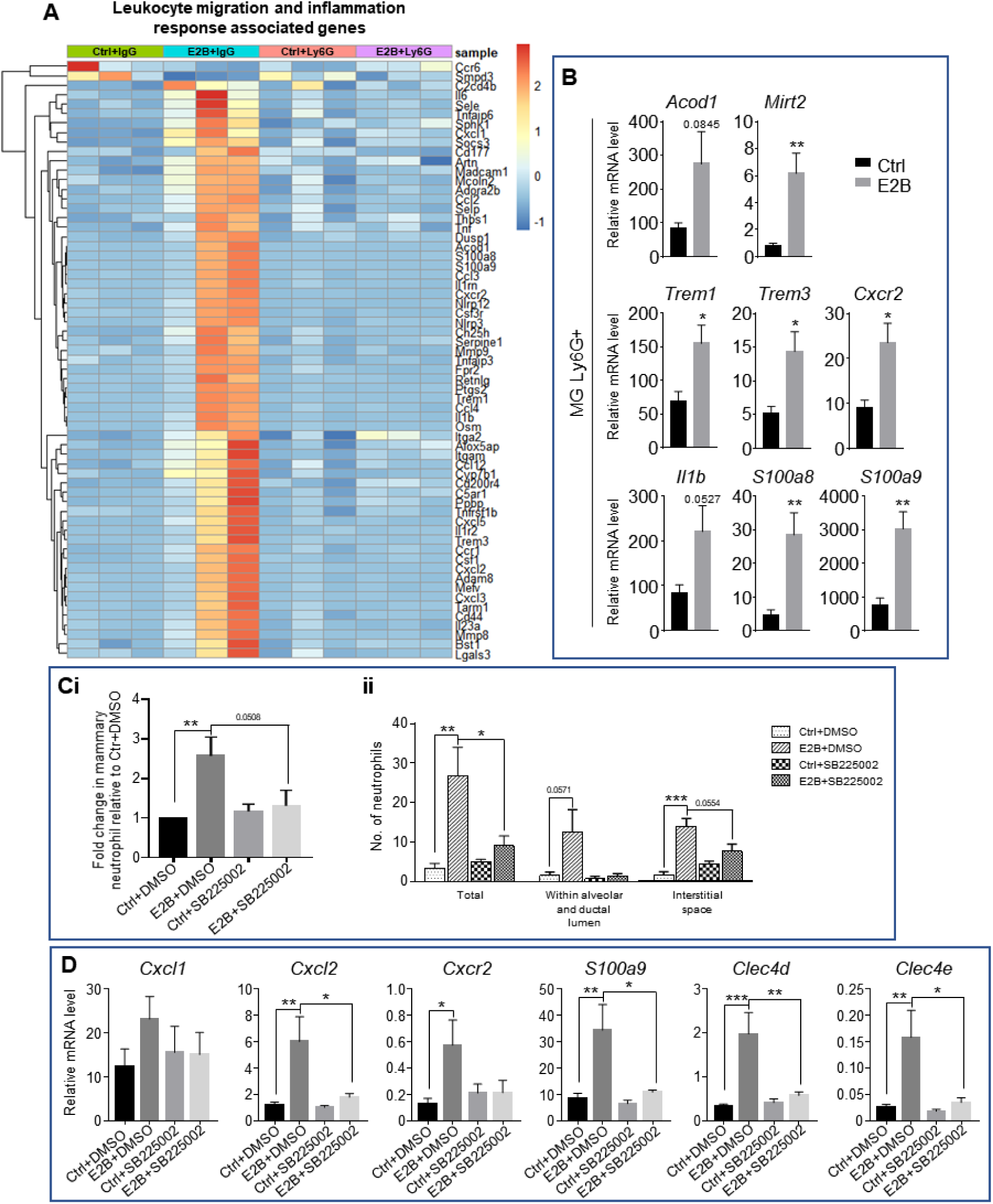
Estrogen-induced *Cxcr2* signalling in neutrophils plays a key role in neutrophil infiltration into the involuting mammary gland. A, Heatmap representation of estrogen-regulated genes associated to leukocyte migration and inflammation in neutrophils (≥ 3 and ≤-3-fold); Experiment is conducted according to the description in Fig. 2. B, qPCR analysis of estrogen-regulated expression of *Acod1*, *Mirt2*, *Trem1*, *Trem3*, *S100a9*, *S100a8*, *Cxcr2*, and *Il1b* relative to *Gapdh* in isolated mammary neutrophils following treatment with or without E2B for 24h (Ctrl n=5, E2B n=5). C-D, E2B-induced *Cxcr2* in neutrophils is critical for E2B-induced neutrophil infiltration. Mice at INV D1 were treated with Ctrl or E2B in the absence or presence of CXCR2 inhibitor SB225002 for 48h; C, SB225002 reduces E2B-induced mammary neutrophil (CD45+ CD11b+ Ly6G+) by flow cytometry analysis (Ci) (Ctrl+DMSO n=7, E2B+DMSO n=7, Ctrl+SB225002 n=7, E2B+SB225002 n=6), and the number of infiltrated neutrophils in 20mm^2^ of mammary sections (Cii) (Ctrl+DMSO n=4, E2B+DMSO n=3, Ctrl+SB225002 n=3, E2B+SB225002 n=3); D, SB225002 reduces E2B-induced *Cxcl2*, *Cxcr2*, *S100a9*, *Clec4d*, and *Clec4e* expression in the involuting gland (Ctrl+DMSO n=7, E2B+DMSO n=6, Ctrl+SB225002 n=7, E2B+SB225002 n=5). Data represented as mean ± SEM.

Next, the study investigated the mechanism of estrogen-induced neutrophils infiltration. S100A8, S100A9, CXCL1, and CXCL2 are known neutrophil chemoattractants that promote neutrophil migration during inflammation [23–25]. S100A8 and S100A9 are small calcium-binding proteins that activate calcium-dependent signalling through receptor for advanced glycation endproducts (RAGE) or toll-like receptor 4 (TLR4). Paquinimod (PAQ) is a derivative of quinoline-3-carboxamide that has been shown to inhibit S100A9 activity through binding with S100A9 [26, 27]. The binding inhibits S100A9 dimerization or the formation of heterodimer with S100A8, thereby preventing the activation of RAGE or TLR4. To test the role of S100A9 in estrogen-induced neutrophil infiltration, PAQ was synthesized following the reported method [28] and tested whether it inhibits E2B-induced neutrophil infiltration by treating OVX mice with E2B or E2B+PAQ at INV D1 for 48h in a pilot experiment. Flow cytometry analysis (Suppl. Fig. 3A) showed that E2B+PAQ treatment resulted in an increase in mammary neutrophils (CD45+ CD11b+ Ly6G+, p=0.0238) and macrophages (CD45+ CD11b+ Ly6C^hi^, p=0.0242) when compared to E2B treated samples. Consistently, qPCR analysis of the treated mammary tissues also revealed a significant increase in pro-inflammatory markers such as *Cxcl2* (p=0.0124), *S100a8* (p=0.0233), *S100a9* (p=0.0161), and C-type lectin domain family 4, member e (*Clec4e*; p=0.0186) (Suppl. Fig. 3C). It is not clear if PAQ has a yet to be characterized functional property that induces neutrophil infiltration, but these results seem to suggest that S100A8/A9 were not significantly involved in the E2B-induced neutrophil infiltration.

Subsequently, we explored the involvement of CXCR2 signalling in E2B-induced neutrophil recruitment because both *Cxcr2* and its ligands *Cxcl1* and *Cxcl2* were both significantly up-regulated by E2B in neutrophils. OVX mice were treated with Ctrl or E2B at INV D1 in the presence of CXCR2 antagonist SB225002 [29], or vehicle control DMSO for 48h. To take into consideration the large variations between experiments, the percentages of infiltrated mammary neutrophils (CD45+ CD11b+ Ly6G+) were presented as fold change over the Ctrl+DMSO group. As shown in Fig. 3Ci, E2B treatment alone without the antagonist (E2B+DMSO) lead to an expected 1.57-fold increase (p=0.0082) in mammary neutrophils as compared to the Ctrl+DMSO. E2B+SB225002 treatment caused a 1.26-fold reduction in mammary neutrophils when compared to the E2B+DMSO group (p=0.0508). Quantification of the number of infiltrated mammary neutrophils within a 20mm^2^ area of stained mammary tissue also showed a significant reduction of neutrophil infiltration within mammary tissue (Fig. 3Cii, p=0.0369). qPCR analysis of the mammary tissue samples showed that the E2B-induced pro-inflammatory markers such as *Cxcl2*, *S100a9*, *Clec4d*, and *Clec4e* were all significantly reduced with E2B+SB225002 treatment compared with E2B+DMSO treatment (Fig. 3D, *Cxcl2*, p=0.0279; *S100a9*, p=0.0199; *Clec4d*, p=0.0051; *Clec4e*, p=0.0183). This is consistent with SB225002-induced inhibition of neutrophil infiltration because these genes are estrogen-induced in neutrophils. Collectively, the data showed that the E2B-induced activation of CXCL2-CXCR2 axis in neutrophils plays a pivotal role in estrogen-induced neutrophil recruitment.

### Estrogen-induced adipocyte repopulation is associated with the induction of adipogenic and tissue remodelling genes through neutrophils

Mammary adipocytes regress during pregnancy and lactation as the mammary epithelial proliferate and differentiate to fill up the mammary fat pad. Adipocytes expansion or repopulation is a hallmark of post-lactational mammary involution. Fig. 1 shows that estrogen significantly induces adipocyte repopulation during mammary involution, and neutrophil depletion attenuates it. RNA-Seq data were further scrutinized and E2B-induced genes associated with fat cell differentiation were identified (Fig. 4A). Intriguingly, E2B down-regulated the expression of *Adipoq*, *Pparg*, and *Lpl* in the IgG group, all of which are positive regulators of adipogenesis. However, E2B also up-regulated some of the upstream regulators of adipogenesis, such as early growth response 2 (*Egr2*; p=0.0001), adipogenin (*Adig*; p=0.0372), CCAAT/enhancer binding protein (C/EBP), beta (*Cebpb*; p=0.0054), and CCAAT/enhancer binding protein (C/EBP), delta (*Cebpd*; p=0.0834), which were validated by qPCR analysis (Fig. 4B), except for *Cebpd* up-regulation which did not achieve statistical significance. In contrast, neutrophil depletion abolished E2B-induced up-regulation of these genes (*Egr2*, p=0.0798; *Adig*, p=0.1542; *Cebpb*, p=0.7571).

**Figure 4.**
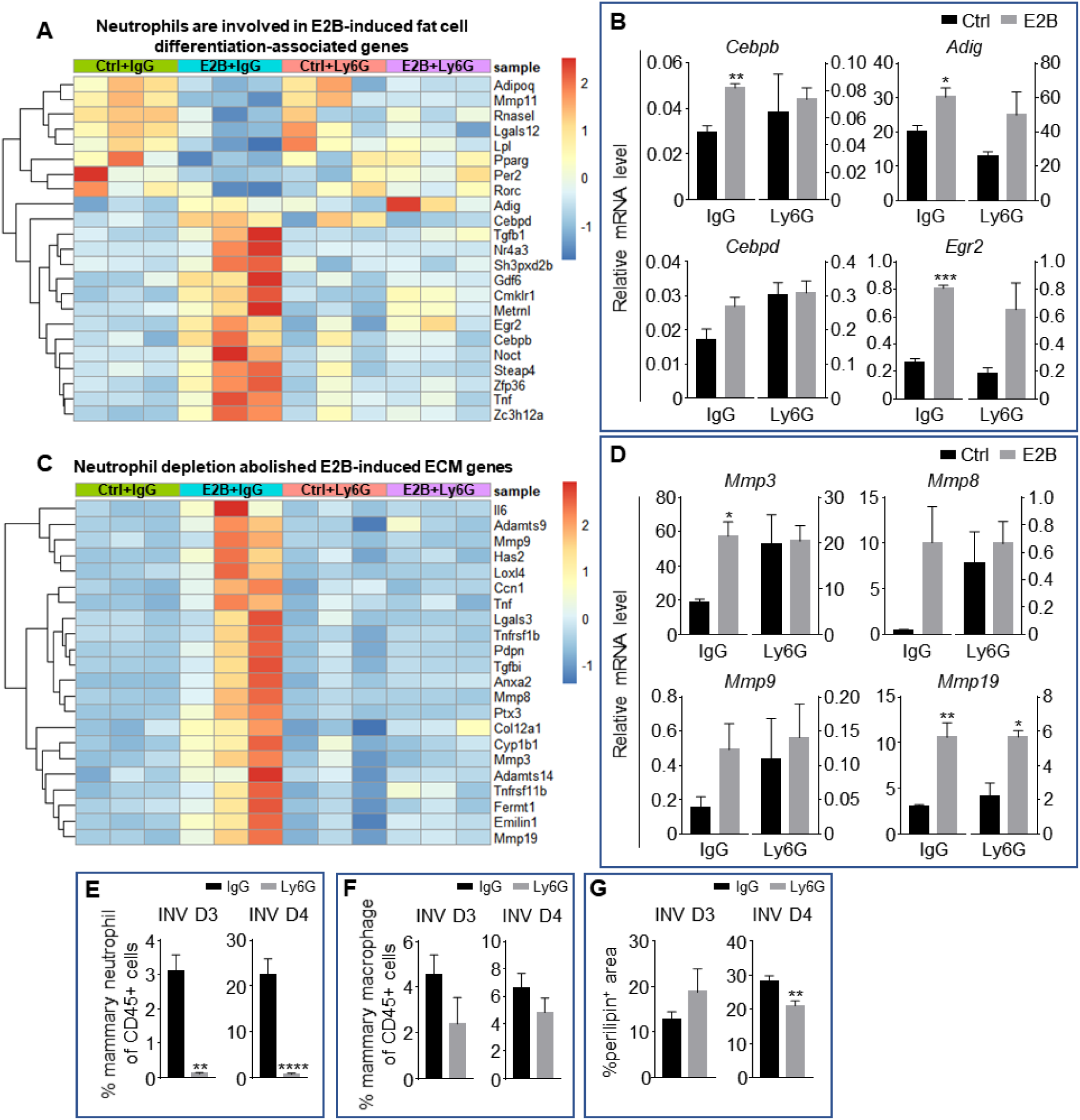
Estrogen-induced adipocyte repopulation is associated with induction of adipogenic and tissue remodelling genes in neutrophils. A, Heatmap representation of genes associated to fat cell differentiation identified from the GO over-representation analysis (≥ 1.5 and ≤ −1.5-fold); Experiment is conducted according to the description in Fig. 2; B, Gene expression of adipogenesis genes *Adig*, *Egr2*, *Cebpb*, and *Cebpd* relative to *36b4* by qPCR analysis. C, Heatmap representation of genes associated to extracellular matrix organization identified from the GO over-representation analysis (≥ 2 and ≤ −2-fold); Section of heatmap replotted from Suppl. Fig. 4; Experiment is conducted according to the description in Fig. 2; D, Gene expression of tissue remodelling genes *Mmp3*, *Mmp8*, *Mmp9*, and *Mmp19* relative to *36b4* by qPCR analysis (Ctrl+IgG n=3, Ctrl+Ly6G n=3, E2B+IgG n=3, E2B+Ly6G n=3). E, Non-OVX mice were treated daily with either anti-Ly6G antibody (Ly6G) or isotype control (IgG) at 24h post-weaning (INV D1); Flow cytometry analysis of mammary neutrophils (CD45+ CD11b+ Gr1^hi^) from mice treated with IgG or Ly6G; F, Flow cytometry analysis of mammary macrophages (CD45+ CD11b+ Ly6C^hi^) from mice treated with IgG or Ly6G; G, Quantification of percentage of perilipin stained area at INV D3 (IgG n=3, Ly6G n=4) and INV D4 (IgG n=10, Ly6G n=9). Data represented as mean ± SEM.

ECM remodelling also plays a key role in adipocyte repopulation during mammary involution [30]. GO over-presentation analysis showed 43 ECM organization associated genes with fold regulation ≥ 2-fold (Suppl. Fig. 4). Six of the down-regulated genes (e.g. *Ecm2*, *Mmp11*, *Col9a1*) were not affected by neutrophil depletion. On the other hand, 22 of the 37 ECM-related genes induced by E2B were reduced with neutrophil depletion (Fig. 4C). Since matrix metallopeptidases (MMPs) are strongly induced in obese adipose tissue and modulate adipocytes differentiation [31], the expression of *Mmp3*, *8*, *9*, and *19* were validated by qPCR. *Mmp3* (p=0.0107) and *19* (p=0.0098) were significantly up-regulated by E2B in the IgG group, and the up-regulation of *Mmp3* was abolished or attenuated with neutrophil depletion (p=0.9353) while *Mmp19* was not affected (p=0.0167) (Fig. 4D). Although the expression level of *Mmp8* was approximately 20 times higher with E2B treatment when compared to Ctrl in the IgG group, it was not statistically significant (Fig. 4D, p=0.0717). This was likely due to considerable variation in neutrophil numbers in these samples as the expression of MMP8 was reported to occur mainly in neutrophils [32]. On the other hand, no significant difference with E2B treatment was observed in *Mmp9* (IgG, p=0.1083; Ly6G, p=0.7036). It should be noted that cytokines *Il6* and *Tnf* were also induced by E2B but both have been reported to inhibit adipogenesis [33, 34]. Hence E2B induced both pro- and anti-adipogenic factors, and the outcome of estrogen-induced adipocytes repopulation could be the result of a balanced act of these genes.

Since adipogenesis genes induced by estrogen appears to be mediated through the estrogen-regulated neutrophil activity, we determined the effect of neutrophil depletion by the Ly6G antibody on adipocytes repopulation in the intact (non-OVX) mice during mammary involution. Mice were given daily injection from INV D1 and mammary gland collected for immunostaining of perilipin at INV D3 (72h post-weaning) showed no effect of the Ly6G antibody (Fig. 4G, p=0.3737 and Suppl. Fig. 5). However, there was a significant reduction in adipocytes repopulation with neutrophil depletion at INV D4 (Fig. 4G, p=0.0077, and Suppl. Fig. 5). The data seems to suggest that neutrophils are involved in adipocytes repopulation during mammary involution normally but the effect is limited to a specific time window when the levels of neutrophil infiltration is high [9].

### Estrogen accelerates lysosomal-mediated programmed cell death during involution

Mammary epithelial cells undergo LM-PCD during the early stage of involution as lysosomes in the mammary cells undergo membrane permeabilization, releasing lysosomal proteases into the cytosol, triggering apoptosis independent of executioner caspases such as caspase 3, 6, and 7 [35–37]. Signal transducer and activator of transcription 3 (STAT3) has been shown to be a regulator for LM-PCD during involution by inducing the expression of lysosomal proteases cathepsin B (CTSB) and L (CTSL), while down-regulating their endogenous inhibitor Spi2A [36].

Despite its well-known function in stimulating mammary growth, estrogen significantly enhanced cell death during the acute phase of mammary involution (Fig. 1A and 1B, p<0.05). Since both RNA-Seq analysis and qPCR demonstrated E2B up-regulation of *Ctsb* expression (Fig. 5A, IgG, p=0.0163; Ly6G, p=0.0182, and Suppl. Fig. 6), we evaluated the protein levels of cathepsins. Indeed, the pro and active form (sc, single-chain) of CTSB were significantly up-regulated (pro-CTSB, p=0.0358; sc-CTSB, p=0.0101) in the E2B-treated mammary gland. Furthermore, the sc-form of cathepsin D (CTSD) and CTSL were also significantly up-regulated by E2B (Fig. 5B, sc-CTSD, p=0.0222; sc-CTSL, p=0.0385), although the levels of pro-CTSD and CTSL were unchanged. Since E2B had no effect on the expression of *Ctsd* and *Ctsl* (data not shown), the increase of the sc-form of CTSD and CTSL can be explained by the up-regulation of CTSB which catalyses the removal of the N-terminal propeptide from itself and from CTSD and CTSL, leading to an increase in active forms of cathepsins B, D, and L [38, 39]. Consistent with the lack of neutrophil influence on E2B-induced cell death, neutrophil depletion had no effect on E2B-induced increase of active form of CTSB (Fig. 5C).

**Figure 5.**
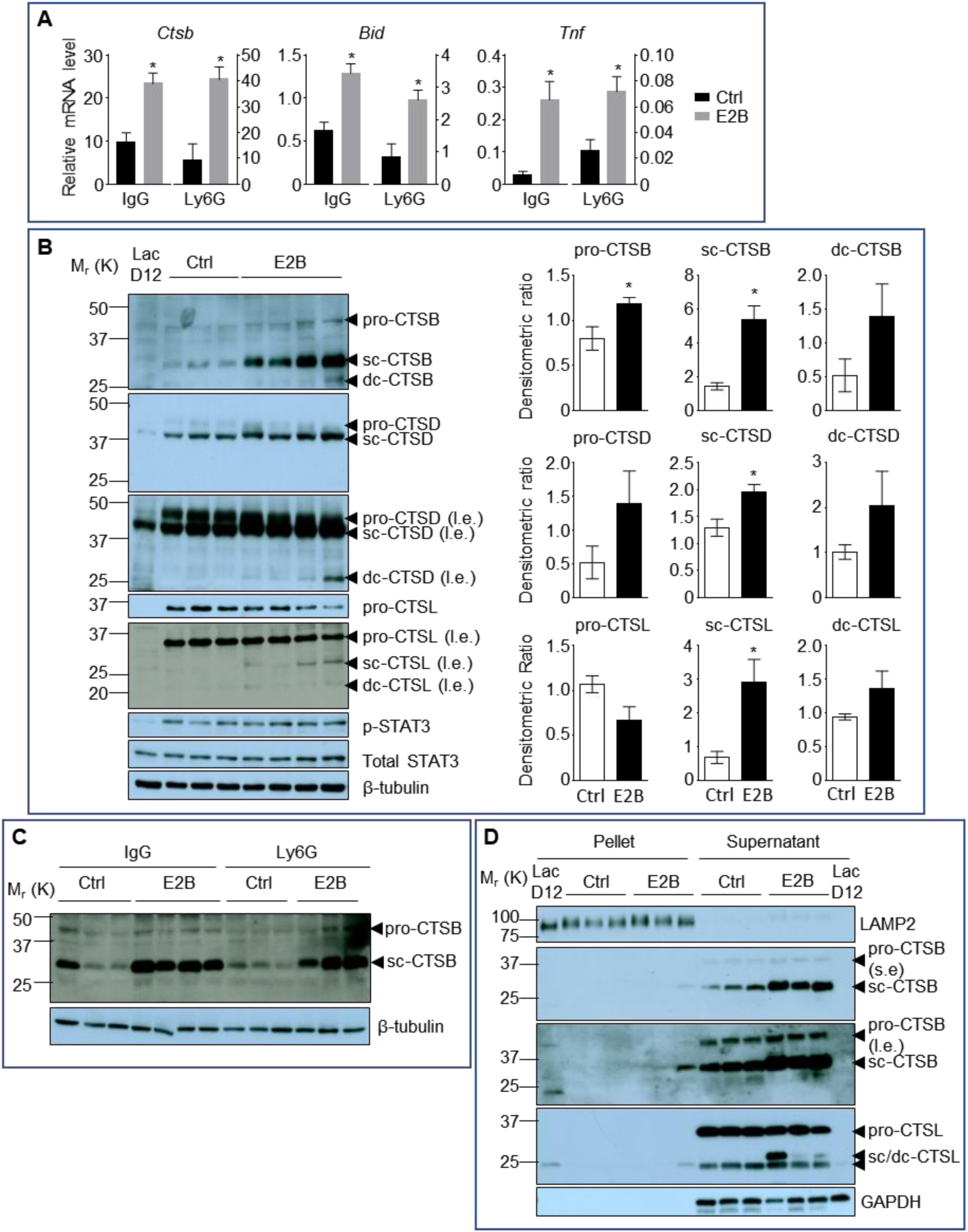
**Estrogen stimulates the activity of lysosomal proteases that are critical for LM-PCD**. A, qPCR validation of E2B-induced expression of *Bid*, *Ctsb*, and *Tnf* relative to *36b4* identified from DESeq2 analysis (Suppl. Fig. 6) (Ctrl+IgG n=3, Ctrl+Ly6G n=3, E2B+IgG n=3, E2B+Ly6G n=3). Mice on INV D1 were treated with Ctrl or E2B for 48h before MG were collected for analysis; B, Western blots of cathepsin B, D and L proteins (sc, single-chain; dc, heavy chain of the double-chain form) in mammary tissue of 48h treatment (Ctrl n=3, E2B n=4). C, Western blotting analysis shows that depletion of neutrophils did not affect estrogen-induced increase of single-chain (sc) and double-chain (dc) forms of CTSB (Ctrl+IgG n=3, E2B+IgG n=4, Ctrl+Ly6G n=3, E2B+Ly6G n=3). D, Effect of E2B on protein levels of lysosomal and cytosolic CTSB and CTSL proteins after subcellular fractionation. LAMP2 is used as a lysosomal marker (s.e, short exposure; l.e., long exposure) (Ctrl n=3, E2B n=3). Data are presented as Mean ± SEM.

E2B also significantly induced the expression of tumour necrosis factor (*Tnf*) (Fig. 5A, IgG, p=0.0161; Ly6G, p=0.0343, and Suppl. Fig. 6), which is a known upstream activator of STAT3. However, E2B did not affect the levels of phosphorylated and total STAT3 (Fig. 5B). This suggests that the up-regulation of *Ctsb* expression by E2B is a direct event independent of STAT3 activation. Furthermore, E2B also up-regulated the expression of BH3 interacting domain death agonist (*Bid*), independent of neutrophils (Fig. 5A, IgG, p=0.0119; Ly6G, p=0.0248, and Suppl. Fig. 6). *Bid* is a well-established pro-apoptotic marker, and its overexpression has been reported to promote cell death [40, 41].

Physiological LM-PCD during mammary involution is associated with the leakage of lysosomal cathepsins into the cytosol. We further tested if estrogen-induced increase of active CTSB and CTSL protein is associated with an increase of cytosolic cathepsins by lysosome and cytosol fractionation. Lysosome-associated membrane protein 2 (LAMP2) was used as a lysosomal marker (Fig. 5D). Consistently, E2B-induced increases of the sc-form of CTSB and CTSL occurred mostly in the cytosolic fraction (Fig. 5D). These findings further support the notion that E2B promotes mammary cell death by increasing the activity of lysosomal proteases CTSB, CTSD, and CTSL under the cellular condition of mammary involution.

Estrogen-induced cell death under physiological condition is hitherto unreported. We hypothesized that pro-inflammatory condition may prime estrogen-induced cell death during the acute phase of mammary involution. As wild-type MCF7 does not express caspase-3 [42], the hypothesis was tested in an *in vitro* model using MCF7-caspase3(+) breast cancer cell line [43]. The idea was to recapitulate estrogen-induced cell death during mammary involution using the pro-inflammatory cytokine TNFα. MCF7-caspase3(+) cells were first treated with TNFα or vehicle control for 1h. This was followed by treatment with 17β-estradiol (E2) or vehicle control for 24h before the cells were stained with propidium iodide (PI) for analysis with flow cytometer. Cells undergoing programmed cell death exhibit increased membrane permeability and hence will stain positive for PI. As shown in Fig. 6A, increasing concentration of TNFα from 2ng/ml to 10ng/ml led to a dose-dependent increase in the percentage of dead cells (p<0.01). As expected, E2B treatment alone did not cause cell death. However, E2B+TNFα treatment significantly enhanced the percentage of dead cells compared to TNFα treatment alone at doses of 5ng/ml (p=0.0104) and 10ng/ml (p=0.0007). Expectedly, TNFα induced increases in p-STAT3 and cleaved poly-ADP ribose polymerase (PARP) protein as compared to the vehicle-treated controls (Fig. 6B). E2B+TNFα increased the levels of p-STAT3 and cleaved PARP significantly compared to TNFα treatment alone (Fig. 6B and 6C, p-STAT3, p=0.0404; cleaved PARP, p<0.0001). These molecular changes are consistent with a greater level of cell death induced by the combined treatment of E2B and TNFα. Together, the *in vitro* data support the notion that pro-inflammatory cytokines such as TNFα primes the death-inducing effect of estrogen. However, the increased p-STAT3 is not associated with increases of active CTSD and CTSB (data not shown), suggesting that estrogen-induced cell death in the presence of TNFα may not be via LM-PCD in the MCF7-caspase3(+) cells.

**Figure 6.**
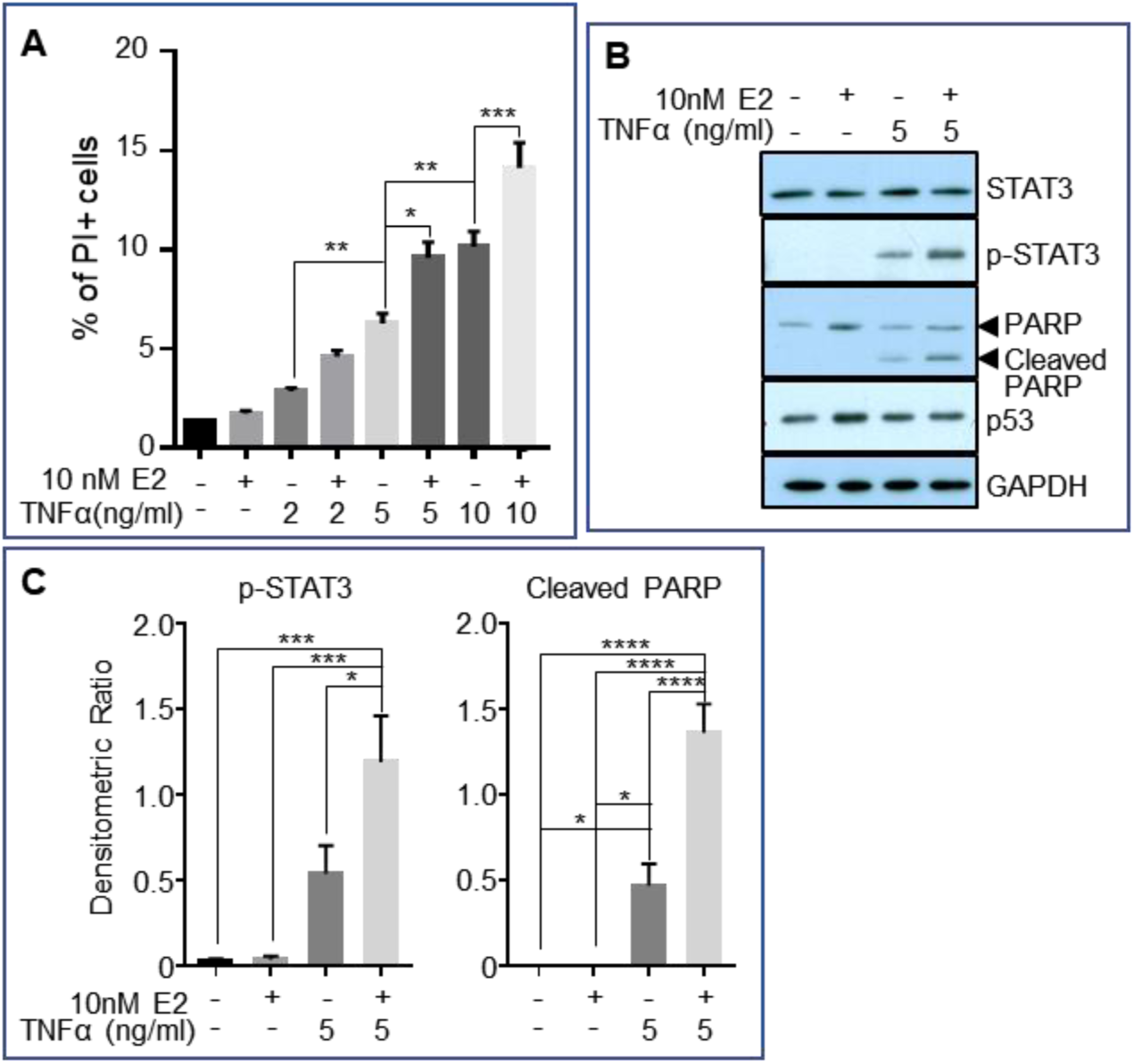
Estrogen accelerates TNFα-induced cell death *in vitro*. MCF7-caspase3(+) cells were treated with either vehicle control (1xPBS) or TNFα of varying concentrations. An hour later, cells were treated with either vehicle control (0.01% ethanol) or 10nM 17β-estradiol (E2) for 24h, after which they were collected for analysis. A, Flow cytometry analysis for the percentage of propodium iodide (PI)-positive cells (dead cells) after treatment (4 independent experiments with triplicates for each group). B, Representative western blotting analysis of various proteins from the treated cells. C, Densitometric analysis of protein expressions normalized against GAPDH (3 independent experiments with duplicates for each group). Data represented as mean ± SEM.

## Estrogen remains a mitogenic hormone during mammary involution

Despite its effect on mammary cell death, estrogen retains its function as a mitogen as indicated by E2B-induced expression of pro-growth genes in the RNA-Seq analysis. The up-regulation of amphiregulin (*Areg*), epiregulin (*Ereg*), and myelocytomatosis oncogene (*c-Myc*) were validated by qPCR in the IgG group (Suppl. Fig. 7, *Areg*, p=0.0448; *Ereg*, p=0.0435; *c-Myc*, p=0.0063) and the E2B’s effects were largely unaffected by neutrophil depletion. These data provide evidence that during post-weaning mammary involution, following the massive cell death events, estrogen promotes the regeneration of the mammary gland.

### The effect of estrogen in age-matched nulliparous mammary tissue is distinct from that undergoing mammary involution

To confirm that the effect of estrogen on neutrophils and on gene regulation during mammary involution does not occur similarly in the post-pubertal mammary gland, OVX nulliparous mice were given isotype IgG the day prior to E2B treatment in order to match the IgG treatment in the involuting mice. 24h later, these animals were given either vehicle control (Ctrl) or E2B for 24h. Flow cytometry analysis of the mammary gland showed that E2B treatment resulted in a 56.33% decrease in mammary neutrophils (CD45+ CD11b+ Gr1^hi^) when compared to Ctrl (Fig. 7A, p=0.0349). E2B had no significant effect on the percentage of mammary macrophage (CD45+ CD11b+ Ly6C^hi^, p=0.2965). qPCR analysis was also performed to investigate the effect of E2B on the expression of pro-inflammatory markers and cell death-related genes. E2B significantly down-regulated *S100a9* (p=0.0095), *Cxcl2* (p=0.0382), and *Cxcr2* (p=0.0236) while had no effect on the expression of *S100a8*, *Clec4d*, *Il1b*, and *Trem3* (Fig. 7C). The expression of *Clec4e*, *Mirt2*, and *Trem1* were also analysed but had no amplification in the qPCR reactions. These observations contrast with that in the involuting mammary gland where the expression of all these pro-inflammatory genes were up-regulated with E2B treatment. As for the cell death-related genes, E2B up-regulated *Ctsb* (p=0.0166), consistent with the understanding that *Ctsb* is an ER target gene [44]. However, E2B had no effect on the expression of *Bid* while *Tnf* displays no amplification in the qPCR reaction due to the low level of expression (Fig. 7D). Hence, unlike its effect during mammary involution, E2B exerts an anti-inflammatory effect on the mammary gland of nulliparous mice.

**Figure 7.**
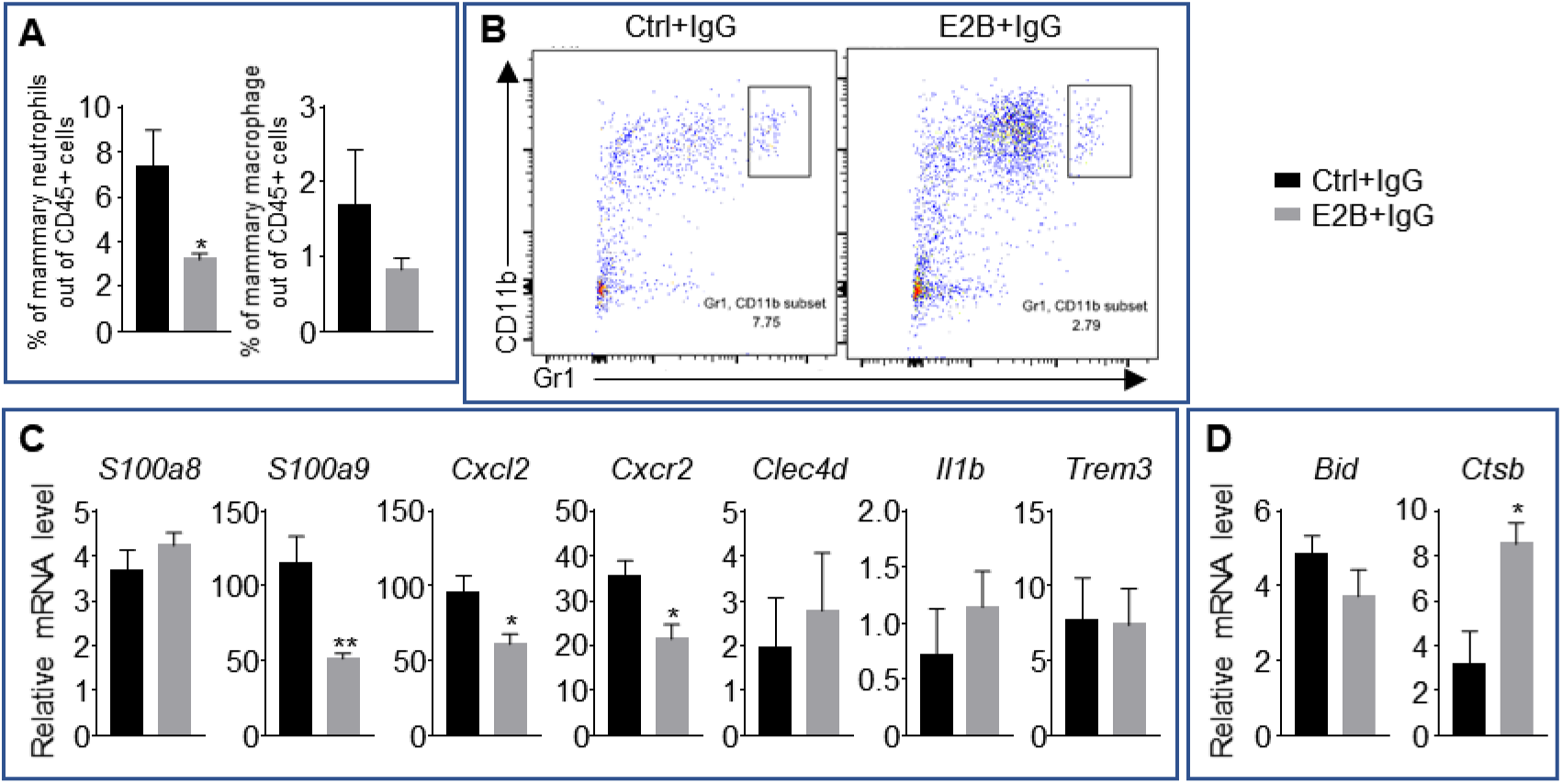
Estrogen regulation of inflammatory and apoptotic genes in nulliparous mammary tissue. OVX nulliparous mice were treated with isotype control (IgG). 24h later, they were treated with either Ctrl or E2B for 24h. A, Flow cytometry analysis of mammary neutrophils (CD45+ CD11b+ Gr1^hi^) and macrophages (CD45+ CD11b+ Ly6C^hi^). B, Representative flow cytometry dot plot for the percentage of neutrophils in the MG. C, Gene expression of pro-inflammatory genes *S100a8*, *S100a9*, *Cxcl2*, *Cxcr2*, *Clec4d*, *Il1b*, and *Trem3* relative to *36b4* by qPCR. D, Gene expression of cell death associated genes *Bid* and *Ctsb* relative to *36b4* by qPCR. Ctrl+IgG n=5, E2B+IgG n=5. All data are presented as mean ± SEM.

## Discussion

This is a comprehensive study of the biological function of estrogen during the acute phase of mammary involution. There are three salient features of the findings. First, estrogen accelerates mammary involution by exacerbating mammary inflammation, programmed mammary cell death, and adipocytes repopulation. These effects were found to be mediated through distinct mechanisms. Second, RNA-Seq analysis revealed the remarkable extent of estrogen regulation of neutrophil activity. Neutrophil depletion eliminated 88% of estrogen-regulated genes in mammary tissue, even though neutrophils account for less than 5% and 1% (in estrogen-treated and control respectively) of the total number of live cells in the mammary tissue tested. Functional analysis showed that neutrophils are the primary mediators of estrogen-induced inflammation, neutrophil infiltration, and adipocyte repopulation. The exceptional impact of estrogen on neutrophil activities would conceivably have a significant influence on the mammary tissue microenvironment. This affirms the pivotal roles of neutrophils in mediating the pro-tumoral effect of estrogen on ER-negative tumour development in mammary tissue during involution [4]. Third, estrogen promotes mammary LM-PCD independent of neutrophils by inducing the expression and activity of lysosomal cathepsins and other pro-apoptotic markers such as *Bid* and *Tnf*. Notably, all these effects of estrogen are unique to the mammary tissue during post-weaning involution, signifying the plasticity of estrogen action depending on the tissue microenvironment.

### Plasticity of estrogen action in neutrophils

In sharp contrast to the effect on neutrophils during mammary involution, estrogen reduced mammary neutrophil infiltration in age-matched nulliparous mice. Estrogen-regulated expression of cytokines such as *S100a9*, *Cxcl2*, and *Cxcr2* in nulliparous mice also occurs in the opposite direction as that in mice undergoing mammary involution. The observation in nulliparous mice is consistent with reports that estrogen inhibits inflammation in obesity-induced mammary inflammation [45], and in *Staphylococcus aureus* infected bovine mammary epithelial cells [46]. To our knowledge, this is the first evidence of the plasticity of estrogen action on neutrophils that is shaped by the tissue microenvironment in an *in vivo* model. This suggests that there are fundamental differences in the cistrome of mammary neutrophils between these two states. It has been shown that TNFα can reshape the genomic action of estrogen in MCF7 cells through redistribution of NF-kB and FOXA1 binding across the genome [47, 48]. The inflammatory microenvironment in mammary tissue during involution may similarly trigger a global shift of ER-cistrome in mammary neutrophils so as to modify the neutrophil response to estrogen.

Neutrophils are known to express ERα, ERβ, and GPER30 [49]. ERα and ERβ are members of the nuclear receptor superfamily of transcription factors, whereas GPER30 is a G protein-coupled membrane receptor. We speculate that ERα plays a major part in regulating the gene expression in neutrophils during mammary involution based on the following evidence. First, the relative *Esr1* (ERα) expression in mammary neutrophils during involution is approximately 40 times more than that in *Esr2* (ERβ) (Suppl. Fig. 8). Second, ERα has been reported to mediate estrogen-induced neutrophil migration in the uterus through ERα phosphorylation at serine 216 [50]. Third, ERα was also reported to mediate the effect of estrogen on myeloid-derived suppressor cells, which are mostly granulocytic cells, in stimulating tumour development in mice model [51]. The cellular factors and the signalling pathway that elicit the epigenetic changes in ERα-cistrome in neutrophils during mammary involution is an interesting area for future study.

### CXCL2-CXCR2 signalling is critical for estrogen-induced neutrophil infiltration

Expectedly, hundreds of estrogen-regulated genes through neutrophils are linked to immune functions such as inflammatory response, chemotaxis, leukocytes adhesion and migration. Using specific CXCR2 antagonist SB225002, this study identified CXCL2-CXCR2 signalling as a major pathway for estrogen to induce neutrophil infiltration. This is consistent with the up-regulation of *Cxcl2* [4], and *Cxcr2* by estrogen in neutrophils (Fig. 3B). CXCR2 has been reported to be important for neutrophil infiltration in several mouse models. In an acute lung injury model, *Cxcr2* gene deletion abolished hyperoxia-induced neutrophil accumulation in the lungs [52]. In studies of reperfusion injury, inhibition of CXCR2 with repertaxin or anti-CXCR2 antibodies led to the reduction of neutrophils accumulation [53–55]. The CXCR2 antagonist SB225002 has also been reported to reduce neutrophil recruitment and pro-inflammatory factor expression in LPS-induced acute lung injury [56]. Although the source of CXCL1 and CXCL2 in those studies were not clear, our data clearly indicate that estrogen-induced *Cxcl2* and *Cxcr2* in neutrophils promote neutrophil recruitment to the mammary tissue.

In addition, CXCL2-CXCR2 signalling likely cooperates with other estrogen-induced chemotactic factors. *Trem1* and *Trem3* are among the top estrogen-regulated genes in neutrophils (Fig. 3A). TREM1 was first identified to be selectively expressed on neutrophils and monocytes [57]. TREM3 is highly homologous to TREM1 and is believed to have overlapping function. TREM1/3-deficient mice displayed impaired neutrophil trans-epithelial infiltration into the lung when challenged with *P. aeruginosa* [58]. TREM1 was also known to amplify inflammation, as TREM1 overactivation with agonistic antibodies following LPS treatment led to the up-regulation of cytokines such as TNFα, MCP-1, and IL8 [57]. It is plausible that *Trem1/3* up-regulation is involved in estrogen-induced neutrophil infiltration.

### Estrogen-stimulated neutrophils play a role in adipocytes repopulation

Estrogen is traditionally known to promote metabolism and inhibit adipogenesis [59]. It has been reported recently that ERα signalling is required for the adipose progenitor identity and the commitment of white fat cell lineage [60]. The present study provides the first evidence that estrogen stimulates adipocytes repopulation during mammary involution, and the effect involves neutrophils based on several lines of evidence. First, neutrophils depletion attenuated estrogen-induced adipocytes repopulation. Second, neutrophil depletion attenuated estrogen-induced expression of *Adig*, *Egr2*, and *Cebpb* which are all known to induce adipocytes differentiation [61–65]. Third, neutrophil depletion in non-ovariectomized mice without estrogen treatment also significantly reduced adipocytes repopulation, although the effect appears to be transient (Fig. 4G). Interestingly, estrogen-induced neutrophil depletion reduced (albeit not significant) infiltrating CD45+ CD11b+ Ly6C^hi^ mammary macrophages in ovariectomized mice (Suppl. Fig. 1B). Mammary macrophages are known to be critical for adipocyte repopulation during mammary involution [66]. It is plausible that estrogen-induced neutrophil recruitment facilitates macrophage infiltration which in turn contributes to adipocytes repopulation.

### Estrogen promotes LM-PCD during mammary involution

The study provides the first evidence that estrogen accelerates cell death when there is ongoing LM-PCD. This is mediated by increased expression of *Ctsb*, and the cytosolic protein levels of active (cleaved) CTSB, CTSD, and CTSL (Fig. 5B). We propose the following model to explain how estrogen-induced *Ctsb* drives LM-PCD. First, increased gene expression and protein levels of CTSB lead to increased levels of activated CTSB due to heightened lysosomal activity in mammary cells with ongoing LM-PCD. The activated CTSB would further increase the cleavage and activation of CTSB, CTSD, and CTSL. CTSB can also enhance the permeability of lysosomal membrane because lysosomal leakage is markedly decreased in cathepsin B-deficient cells in respond to TNFα treatment [67]. Hence, estrogen-induced expression of *Ctsb* triggers a chain of events, culminating in increased LM-PCD. In addition, the estrogen-induced expression of *Tnf* and *Bid* further enhance this process. Although estrogen-induced *Tnf* is not associated with increased activation of STAT3, TNFα has been reported to redistribute the zinc transporter ZnT2 to increase Zn in lysosomes. The high levels of Zn cause lysosomal swelling and cathepsin B release during mammary involution [68, 69]. It is well known that cathepsins cleave BID to tBID and degrade antiapoptotic BCL-2 homologues to execute LM-PCD [70, 71]. Increased *Bid* level would thus reinforce cathepsin-stimulated LM-PCD. Hence, estrogen-induced increases of cytosolic cathepsins together with increase of *Tnf* and *Bid* accelerate LM-PCD during mammary involution.

On the other hand, estrogen induction of *Ctsb* in nulliparous mice (Fig. 7D) without ongoing LM-PCD would not lead to cell death due to the lack of a permissive cellular environment. This notion is supported by the study of estrogen in MCF7-Caspase3(+) cells, in which treatment with TNFα primes estrogen to stimulate cell death. In contrast to the observation in involuting mammary gland in which estrogen did not increase total or pSTAT3, estrogen plus TNFα enhanced STAT3 phosphorylation as compared to TNFα alone (Fig. 6C). However, TNFα did not affect the levels of cathepsins significantly in MCF7-Caspase3(+) cell, and hence it unlikely induced LM-PCD. Nonetheless, the *in vitro* study demonstrates that estrogen can be primed to heighten cell death under a pro-inflammatory stimulus. Estrogen has also been reported to induce apoptosis in experimental endocrine resistance models through inducing endoplasmic reticulum stress and inflammatory response [72]. Thus, estrogen can induce cell death under certain cellular context, but the mechanisms vary.

## Conclusion

In summary, the study reveals distinct novel mechanisms of estrogen action that are unique to the mammary tissue during post-weaning mammary involution (Fig. 8). Estrogen treatment intensely and extensively regulates the activity of neutrophils to elicit inflammation and adipocytes repopulation. The study identified CXCL2-CXCR2 signalling in neutrophils as the main mechanism of estrogen-induced neutrophil recruitment. Additionally, estrogen also exacerbates mammary LM-PCD by inducing the expression of *Ctsb*, *Tnf*, *Bid*, and subsequent lysosomal activation and release of CTSB, CTSD, and CTSL independent of neutrophils. At the same time, estrogen retains its function in inducing the expression of pro-growth genes to facilitate mammary regeneration for the subsequent reproduction. It should also be recognized that programmed cell death is often associated with growth, when the elimination of unwanted cells can benefit tissue remodelling and regeneration [73]. For example, it has been shown in Drosophila eye imaginal discs that apoptotic cells release Spi, the EGF ligand in flies, to promote the proliferation of neighbouring cells [74]. Whether estrogen-induced LM-PCD during mammary involution additionally facilitates mammary regeneration and tumour development is an interesting area for future study.

**Figure 8.**
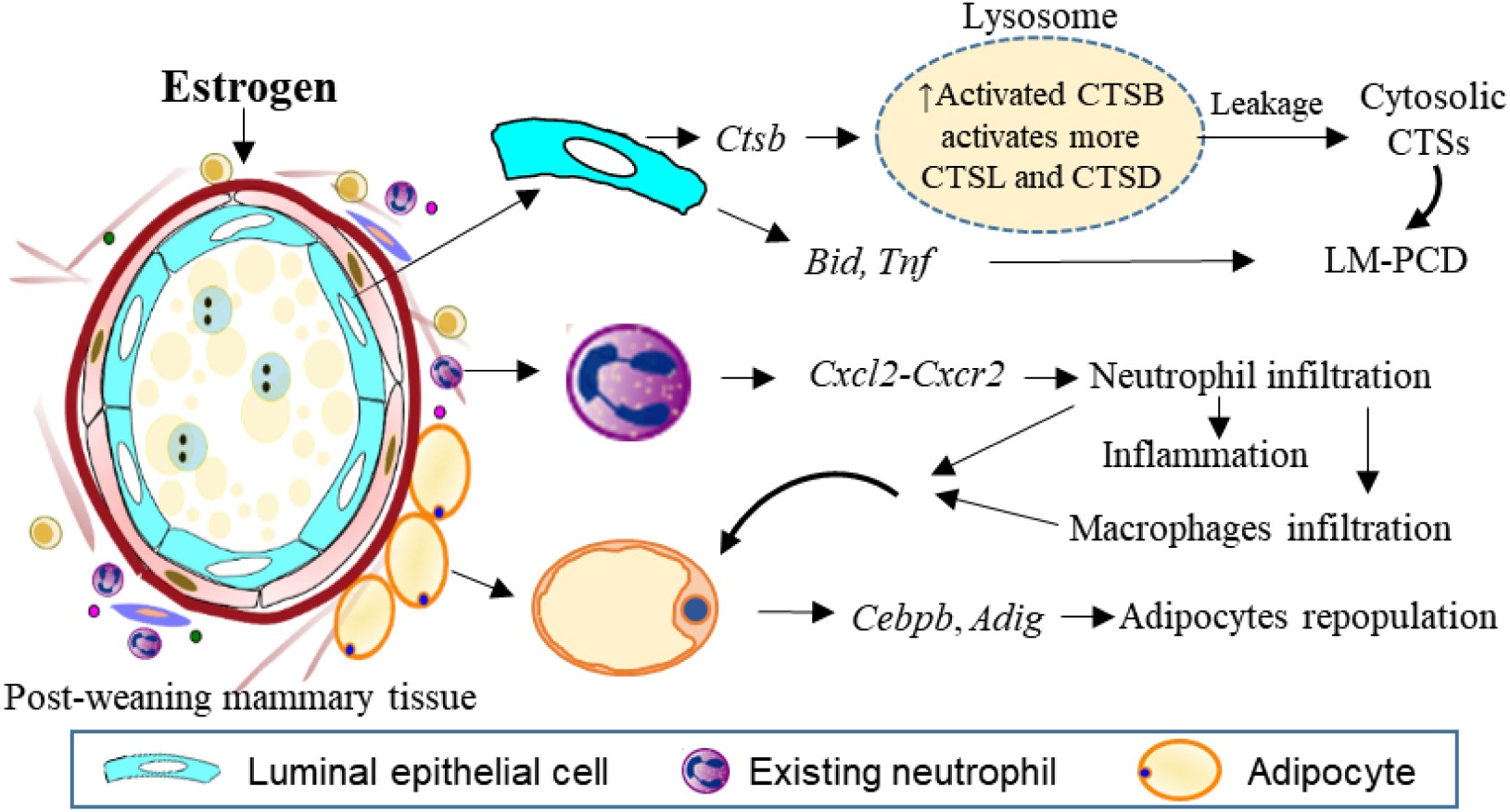
Estrogen exacerbates mammary cell death, inflammation and adipocytes repopulation through distinct mechanisms. Estrogen induces *Ctsb* gene expression in the mammary cells leading to increased pro-CTSB which is cleaved and activated in lysosomes. The increased active sc-CTSB further activates CTSD and CTSL. Increased CTSB activity enhanced lysosomal permeabilization during mammary involution resulting in the leakage of more activated CTS into the cytosol stimulating a heightened LM-PCD. Estrogen treatment also induces expression of *Bid* and *Tnf* gene which are reported to be involved in the induction of LM-PCD. The apoptotic protein BID is cleaved into the active tBID by the activated cytosolic CTSs. TNFα is known to induce LM-PCD via the ZnT2-mediated zinc accumulation in lysosomes, leading to PCD. The study also finds that estrogen stimulates neutrophil infiltration into the involuting mammary gland via the CXCL2/CXCR2 pathway. Meanwhile, estrogen promotes the expression of numerous proinflammatory genes such as *Trem1, Trem3, Il1b, S100a8, S100a9* in neutrophils that heighten mammary inflammation. Furthermore, increased neutrophil infiltration can also recruit macrophage into the involuting gland contributing to the observed estrogen induced adipocyte repopulation. Estrogen-induced expression of genes coding for extracellular matrix remodelling enzymes such as *Mmp19, Mmp3, Mmp8, Ptx3, Col8a2, Has2* further facilitate the adipocyte repopulation during mammary involution.

Both circulating and tissue neutrophils are known to undergo functional and phenotypic changes during cancer development and under other pathological conditions [14, 75]. It appears that neutrophils in the post-weaning, inflammatory involuting mammary tissue have undergone epigenetic changes that are particularly responsive to estrogen in fostering the pro-tumoral microenvironment. It is tempting to speculate that neutrophils in inflammatory breast cancer also undergo similar epigenetic changes that are responsive to the pro-inflammatory effect of estrogen. Further understanding of the epigenetic properties of mammary neutrophils during involution will yield biomarkers that are predictive pro-tumoral neutrophils.

## Materials and methods

### Animal studies

All animal experiments were performed in accordance with the protocol approved by the Nanyang Technological University Institutional Animal Care and Use Committee (NTU-IACUC) (IACUC protocol number: A0306 and A18036). BALB/cAnNTac mice used in the study were housed in specific pathogen-free (SPF) facility under a 12 hours dark/light cycle and provided food and water ad libitum. Female mice at 7 to 8 weeks old were mated and subsequently housed individually prior to giving birth. Bilateral ovariectomy (OVX) was performed on the lactating mice two days post-partum to remove the endogenous source of estrogen. The litter sizes were standardized to 5-6 pups per lactating mouse. Mammary involution was initiated by forced weaning on lactating day 12. Nulliparous mice were handled similarly to the pregnant mice. All mice were randomly distributed into the different treatment groups.

### Drug and estrogen treatment

17β-estradiol-3-benzoate (E2B) (Sigma-Aldrich) dissolved in benzyl alcohol was administrated subcutaneously at 20µg/kg of body weight per day in sesame oil. Control (Ctrl) animals received sesame oil at the corresponding volume per body weight.

CXCR2 inhibitor SB225002 (Sigma-Aldrich) was dissolved in DMSO to a stock concentration of 20mg/ml. Mice were treated twice daily at a dosage of 0.3mg/kg body weight via intraperitoneal injection. Stock concentration of SB225002 was diluted to a working concentration of 0.2mg/ml with 0.25% (v/v) Tween 20 (Bio-Rad) in 1xPhosphate-buffered saline (PBS) (Vivantis). Control mice were treated with DMSO in 0.25% Tween 20 in 1xPBS.

Putative S100A9 inhibitor Paquinimod (PAQ) or ABR-215757 was synthesized (Suppl. Fig. 9) by following the reported protocol [28]. The stock solution was prepared by dissolving PAQ in DMSO to a concentration of 50mg/ml. Mice were treated with a daily dosage of 20mg/kg body weight via intraperitoneal injection of PAQ diluted with 1xPBS to a working concentration of 5mg/ml. Control mice were treated with DMSO in 1xPBS.

### Neutrophil depletion

At 24h post-weaning, involution day 1 (INV D1), mice were administered with either rat IgG2a isotype control (clone 2A3, BioXCell) or rat monoclonal anti-mouse Ly6G antibody (clone 1A8, BioXCell) via intraperitoneal injection. Antibodies were administered at a daily dosage of 20µg in 100µl of 1xPBS. For nulliparous animals, antibody treatment was performed at 24h prior to hormone injection. The efficiency of neutrophil depletion by the neutralizing antibodies was determined by flow cytometry analysis of the blood and the mammary gland.

### RNA-sequencing (RNA-Seq)

OVX female mice were treated with either anti-Ly6G or IgG antibody at INV D1. At INV D2, mice were then treated with either Ctrl or E2B for 24h. Mammary tissues were collected and snap-frozen in liquid nitrogen and stored at −80°C. Total RNA was extracted from the powdered 9^th^ abdominal mammary gland using TRIzol reagent (Life technologies) followed by treatment with DNase I (DNA-free™ DNA Removal Kit, Invitrogen) to remove contaminating DNA according to the manufacturers’ protocol. The total RNA was then sent for library preparation and paired-end sequencing by A*STAR GIS (Agency for Science, Technology and Research, Genome Institute of Singapore) using Illumina HiSeq4000. The Illumina adapter sequence was removed using Trimmgalore. Processed sequences were subsequently mapped to the *Mus musculus* BALB/cJ reference genome (obtained from Ensembl) and counted using the stringtie and featurecount program. Gene annotation files were also obtained from Ensembl.

Processed RNA-Seq data were analysed for differential gene expression using the DESeq2 package with contrast method [21]. Statistically significant differential gene expression was determined by Benjamini-Hochberg adjusted p-value (padj). Volcano plot based on the results of DESeq2 analysis was generated using the plotly package [76]. Venn diagram was plotted using the online software Venny [77]. Pathway analysis of the differentially expressed (DE) gene (padj<0.05) was then conducted using the clusterProfiler package [78] where gene ontology (GO) over-representation analysis was performed. The enriched GO terms obtained from the GO over-representation analysis were removed of redundancy using the ‘simplify’ function which removes highly similar enriched GO terms and keeps only one representative term. The DESeq2, plotly, and clusterProfiler package was run in R using RStudio [79, 80].

### Isolation of mammary neutrophils

Mammary tissues collected from the euthanized mice were minced into small pieces of approximately 1mm. Minced mammary tissues were then digested in phenol-red free Dulbecco’s Modified Eagle Medium (DMEM) (Nacalai Tesque) containing 2mM L-glutamine (GE healthcare), 1mg/ml collagenase (Sigma-Aldrich), and 120 Kunitz DNase I (Sigma-Aldrich) for 1h at 37°C in water bath with agitation. Digestion was inactivated with an equal volume of medium and cells were sieved through a 100µm sieve and centrifuged at 450g for 5min at 4°C. Red blood cells (RBC) lysis was then performed with 1ml NH4Cl buffer (1 volume of 0.17M Tris-HCL and 9 volume of 0.155M NH4Cl). After inactivation of RBC lysis with 10 volume of 1xPBS, cells were pellet and incubated with 120 Kunitz/ml DNase I for 15min at room temperature. Cells were then centrifuged again and removed of supernatant before resuspension in isolation buffer (calcium and magnesium-free 1xPBS, 0.1% (w/v) bovine serum albumin (BSA) (Cell Signaling Technology), 2mM EDTA, pH7.4).

Isolation of mammary neutrophils using Dynabeads® (Thermo Fisher Scientific) from the digested mammary cells were carried out following the manufacturer’s protocol. Briefly, 5 million cells from the mammary tissue digest were incubated with 1µg of biotin-anti-Ly6G antibody (clone 1A8, BioLegend) at 4°C for 10min. After washing, cells were then incubated with 10µl of Dynabeads® for 30min at 4°C with gentle rotation and tilting. Dynabeads®-bound cells were separated from the non-bound cells using a magnetic stand (Milipore) on ice. Isolated Dynabeads® bound neutrophils were washed and then added TRIzol, snap-frozen with liquid nitrogen and stored at −80°C. A small amount of the non-bound cell fraction after washing was used for flow cytometry analysis of cell depletion efficiency. Remaining non-bound cells were also collected in TRIzol, snap-freeze and stored at −80°C. A concurrent separate isolation was also performed with the same above described protocol with no antibody incubation. This was done to obtain a negative control to ensure the specificity of the Dynabeads® isolation process.

### Quantitative PCR (qPCR)

Following the isolation of total RNA with TRIzol, reverse transcription was carried out using qScript cDNA SuperMix (Quantabio) following the manufacturer’s protocol. qPCR was carried out with KAPA SYBR FAST qPCR Master Mix (KAPA Biosystems) on the Quantstudio 6 Flex Real-Time PCR System (Applied Biosystems). qPCR for each target gene was performed in duplicates. For quantitative analysis, the comparative Threshold Cycle (*C*t) method was used, while normalizing to *C*t value of *36b4* or *Gapdh* in the same sample. Relative quantification was performed using the 2^−Δ*C*t^ method [81]. The data are expressed as relative mRNA level in arbitrary values. Primers are listed in Suppl. Table 1.

### Histological analysis

4% paraformaldehyde-fixed mammary tissue samples (4^th^ abdominal mammary gland) were paraffin-embedded and sectioned at 5µm for haematoxylin and eosin (H&E) staining and immunohistochemical (IHC) staining. To quantify epithelial cell death, the number of dying cells shed into the alveolar lumen of the mammary gland was counted. Apoptotic cells were identified by their morphological characteristics as described previously [37]. IHC staining was carried out using the VECTASTAIN^®^ Elite® ABC Kit (Vector laboratories) and perilipin A (1:100, D418 Cell Signaling) or cleaved caspase 3 (1:100, #9661 Cell Signaling) followed by the DAB (3,3’-diaminobenzidine) peroxidase (HRP) substrate kit (with Nickel) (Vector Laboratories). The tissue sections were counterstained with Richard-Allan Scientific^TM^ Signature Series Hematoxylin 2. All histological analysis was performed with images from at least 5 random fields for each sample.

### Subcellular fractionation

Subcellular fractionation of mammary glands was carried out based on the published protocol [36]. Briefly, mammary tissues in liquid nitrogen were powdered and homogenized in a handheld homogenizer in subcellular fractionation buffer (20mM HEPES-KOH, 250mM sucrose, 10mM KCl, 1.5mM MgCl2, 1mM EDTA, 1mM EGTA, 8mM dithiothreitol, 1mM Pefabloc, at pH7.5) and centrifuged at 750g for 10min at 4°C to remove cell nuclei and debris. The supernatant was then spun at 10,000g for 15min at 4°C to pellet organelles. The pellet was washed and re-suspended in subcellular fractionation buffer as lysosomal fraction. Organelles were disrupted by three cycles of freezing and thawing. To collect the cytosolic fraction, the supernatant collected after pelleting organelles was spun at 100,000g for 1h at 4°C to remove microsomes. Protein concentration was determined by the Bradford protein assay (Bio-rad).

### Flow cytometry analysis

Blood cells collected via cardiac puncture were lysed of RBC via incubation with NH4Cl buffer for 15min at room temperature with gentle tilting. RBC lysis was inactivated with equal volume of 1xPBS followed by centrifugation at 450g for 5min at 4°C and resuspended in 1xPBS. Mammary tissues were digested as previously described until after the RBC lysis step in which mammary cells were subsequently reconstituted in 1xPBS. Staining of cells was carried out with a cocktail of primary antibodies from either BioLegend or eBioscience comprising of APC-Cy7-anti-CD45 (clone 30-F11), FITC-anti-Ly6C (clone HK1.4), PE-anti-Ly6G (clone 1A8), Biotin-anti-Gr1 (clone RB6-8C5), and BV605-anti-CD11b (clone M1/70). Secondary antibody staining was performed with Alexa Fluor® 647-streptavidin antibody (BioLegend). Dead cells were stained with eFluor 450-fixable viability dye (eBioscience) before fixation with fixative buffer (BioLegend).

### Protein collection and western blotting

Proteins were isolated from cells or tissues by adding cold lysis buffer containing 50mM HEPES (pH7.5), 150mM NaCl, 100mM NaF, 1mM PMSF, 1% (v/v) Triton X-100, and a cocktail of proteinase inhibitors (2ug/ml aprotinin, 5ug/ml leupeptin, 1mM Na3VO4, and 5ug/ml pepstatin A). After lysis, cells were centrifuged at 13800rpm for 15min at 4°C and the supernatant collected. Protein concentration was determined with the BCA protein assay kit (Thermo Fisher Scientific) following the manufacturer’s protocol. The collected protein lysate supernatant was added 5x Laemmli sample buffer and stored at −80°C. Protein samples collected were analysed with western blotting. The protein band of interest was subsequently quantitated (normalized to the reference protein band) using quantity one software (Bio-rad). Antibodies used for western blot analysis are listed in Suppl. Table 2.

### *In vitro* studies of estrogen-induced cell death

MCF7-caspase3(+) cells were used in order to observe TNFα-induced apoptosis [43]. Cells were maintained in DMEM containing 2mM L-glutamine and 7.5% fetal calf serum (FCS) (HyClone) and kept at 37°C in a humidified 5% carbon dioxide and 95% air atmosphere.

For estrogen treatment, phenol-red free DMEM supplemented with 2mM L-glutamine and 5% DCC-FCS (fetal calf serum treated with dextran-coated charcoal) was used. Treatment of FCS with dextran-coated charcoal was performed to remove steroid hormones present in the FCS. MCF7-caspase3(+) cells were plated at 150k in 6-well plate (Corning) with the DCC-FCS supplemented medium. After 48h, the medium was replaced, and cells were treated with either TNFα (ProSpec) diluted in 1xPBS or vehicle control. One hour after TNFα treatment, cells were then treated with 10nM 17β-estradiol (E2) diluted in 100% ethanol or vehicle control. MCF7-caspase3(+) cells were harvested for analysis 24h after E2 treatment. Both floating dead cells and the adherent live cells were collected. Harvested cells were resuspended in 1xPBS and stained with 0.1μg PI per 100ul of 1xPBS for 15min at room temperature. After staining, 400μl of 1xPBS were added and cells were immediately analysed with the flow cytometer.

### Statistical analysis

Graphs were plotted using the mean value with the standard error of the mean (SEM). When comparing 2 groups, statistical significance was determined using a two-tailed unpaired student’s t-test. When comparing between more than 2 groups, one-way ANOVA followed by post-hoc turkey test was performed. All statistical analysis was performed using the GraphPad Prism 7 software. p-value: <0.05 (*), <0.01 (**), <0.001 (***), <0.0001 (****).

## Acknowledgements

This research is funded the Ministry of Education of Singapore. Academic Research Fund Tier I, MOE2017-T1-002-08. We thank Drs. Natasa Bajalovic, Amanda Woo and Mr. Lee Shi Hao for their technical assistance.

## Competing interest

The authors declare that they have no competing interests.

## Supplementary figures

**Supplementary Figure 1.**
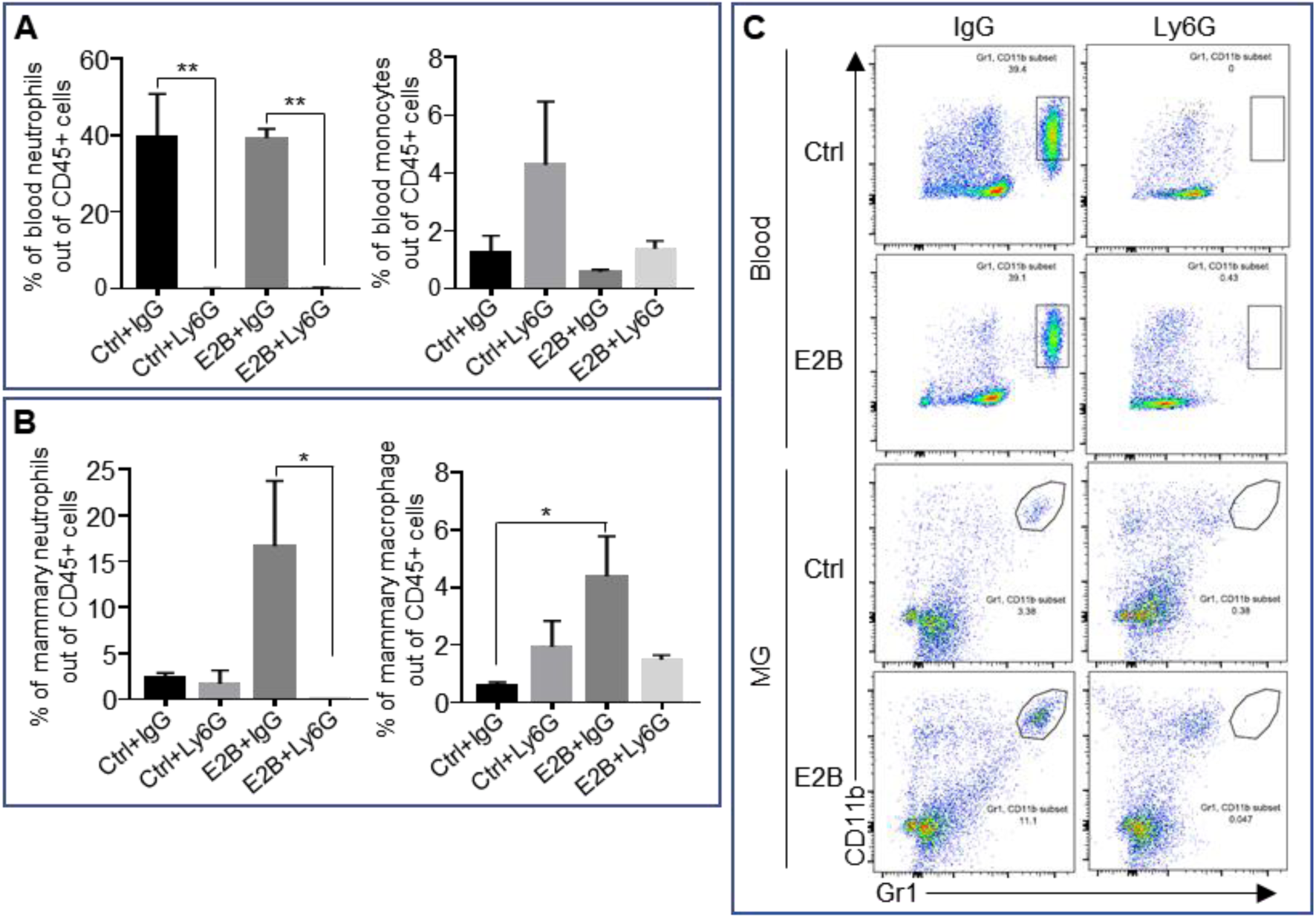
Depletion efficiency of neutrophils with anti-neutrophil antibody Ly6G. Mice at INV D1 was treated with anti-Ly6G antibody (Ly6G) or isotype control (IgG). 24h later, they were treated with vehicle control (Ctrl) or E2B for 24h. Flow cytometry analysis were performed on total blood cells and from digested mammary gland (MG) tissue after red blood cells lysis. A, Ly6G treatment significantly reduces circulating blood neutrophils by more than 90% while not affecting blood monocytes; Percentage of blood neutrophils (CD45+ CD11b+ Gr1^hi^) and monocytes (CD45+ CD11b+ Ly6C^hi^) out of live CD45+ population. B, E2B treatment in mice given IgG increased the percentage of mammary neutrophils by 6 folds and this effect was abolished by neutrophil depletion with Ly6G. Mammary macrophages was also increased significantly by E2B treatment and was attenuated with Ly6G but to a non-statistically significant level; Percentage of mammary neutrophils (CD45+ CD11b+ Gr1^hi^) and macrophages (CD45+ CD11b+ Ly6C^hi^) out of live CD45+ population. Ctrl+IgG n=3, Ctrl+Ly6G n=3, E2B+IgG n=3, E2B+Ly6G n=3. C, Representative flow cytometry dot plot for the neutrophils in the blood and MG. Data represented as mean ± SEM.

**Supplementary Figure 2.**
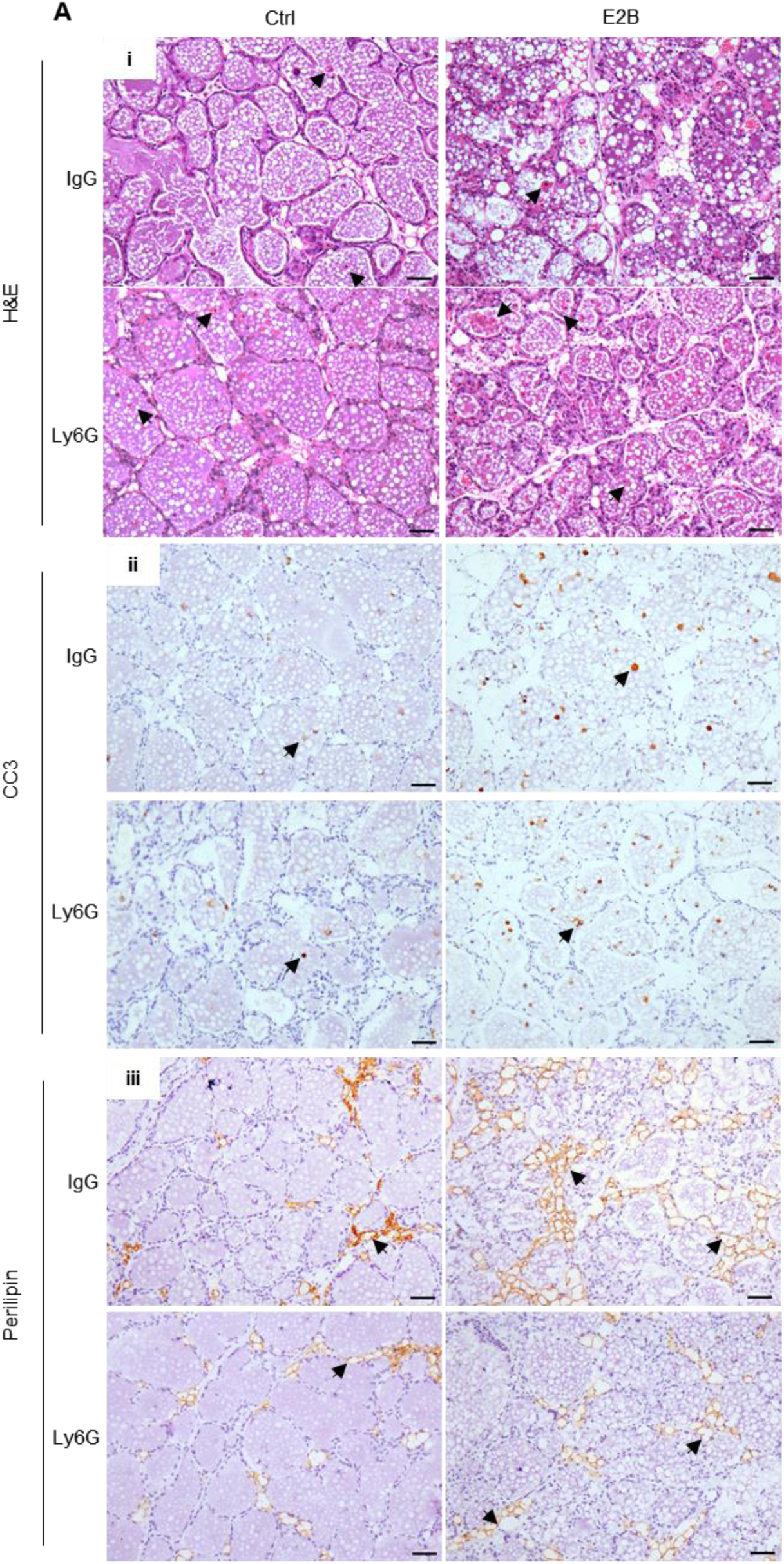
Effect of neutrophil depletion on estrogen-induced cell death and adipocytes repopulation. Mice at INV D1 were administered with isotype control (IgG) or anti-neutrophil antibody (Ly6G) and treated with Ctrl or E2B for 48h. A, Neutrophil depletion attenuates E2B-induced adipocyte repopulation but did not affect the E2B-stimulated cell death in involuting mammary gland; (i) H&E stained mammary tissue sections; shed cells with hyper-condensed nuclei are indicated by arrows. (ii) IHC of cleaved caspase-3 (CC3); arrows indicate CC3^+^ cells. (iii) Perilipin IHC; arrows indicate perilipin^+^ adipocytes. Scale bars: 50µm.

**Supplementary Figure 3.**
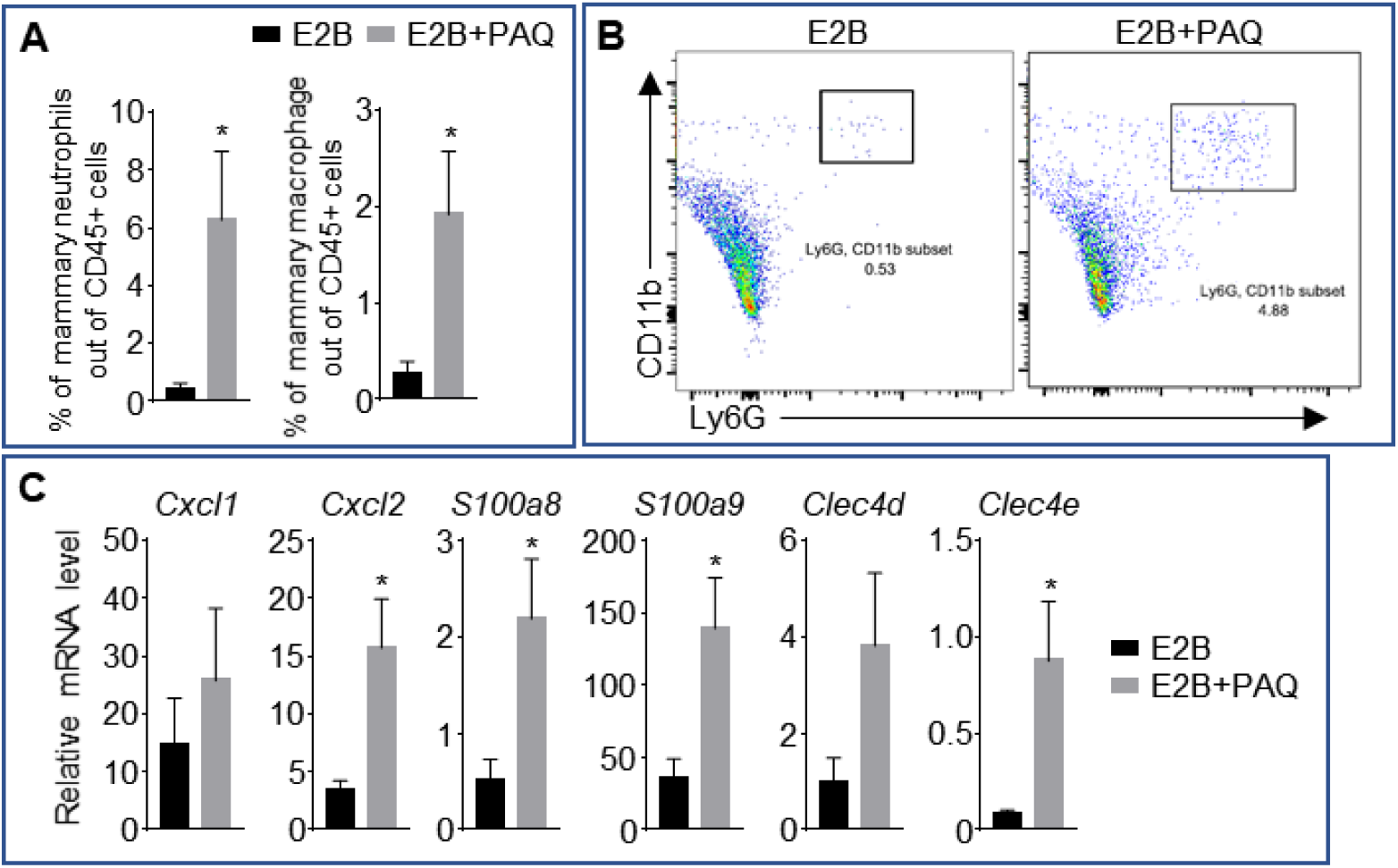
Putative S100A9 inhibitor Paquinimod promotes neutrophil infiltration. Mice were treated at INV D1 with either E2B or E2B+PAQ for 48h. A, Treatment with E2B and PAQ significantly induces neutrophil and macrophage infiltration into the involuting MG as compared to E2B-treated; Flow cytometry analysis of mammary neutrophils (CD45+ CD11b+ Ly6G+) and macrophages (CD45+ CD11b+ Ly6C^hi^) out of live CD45+ cell population. B, Representative flow cytometry dot plot for the percentage of neutrophils in the MG. C, Treatment with E2B and PAQ increases the expression of some inflammatory genes as compared to E2B-treated; Gene expression of inflammatory genes *Cxcl1*, *Cxcl2*, *S100a8*, *S100a9*, *Clec4d*, and *Clec4e* relative to *36b4* by qPCR analysis. E2B n=6, E2B+PAQ n=5. Data represented as mean ± SEM.

**Supplementary Figure 4.**
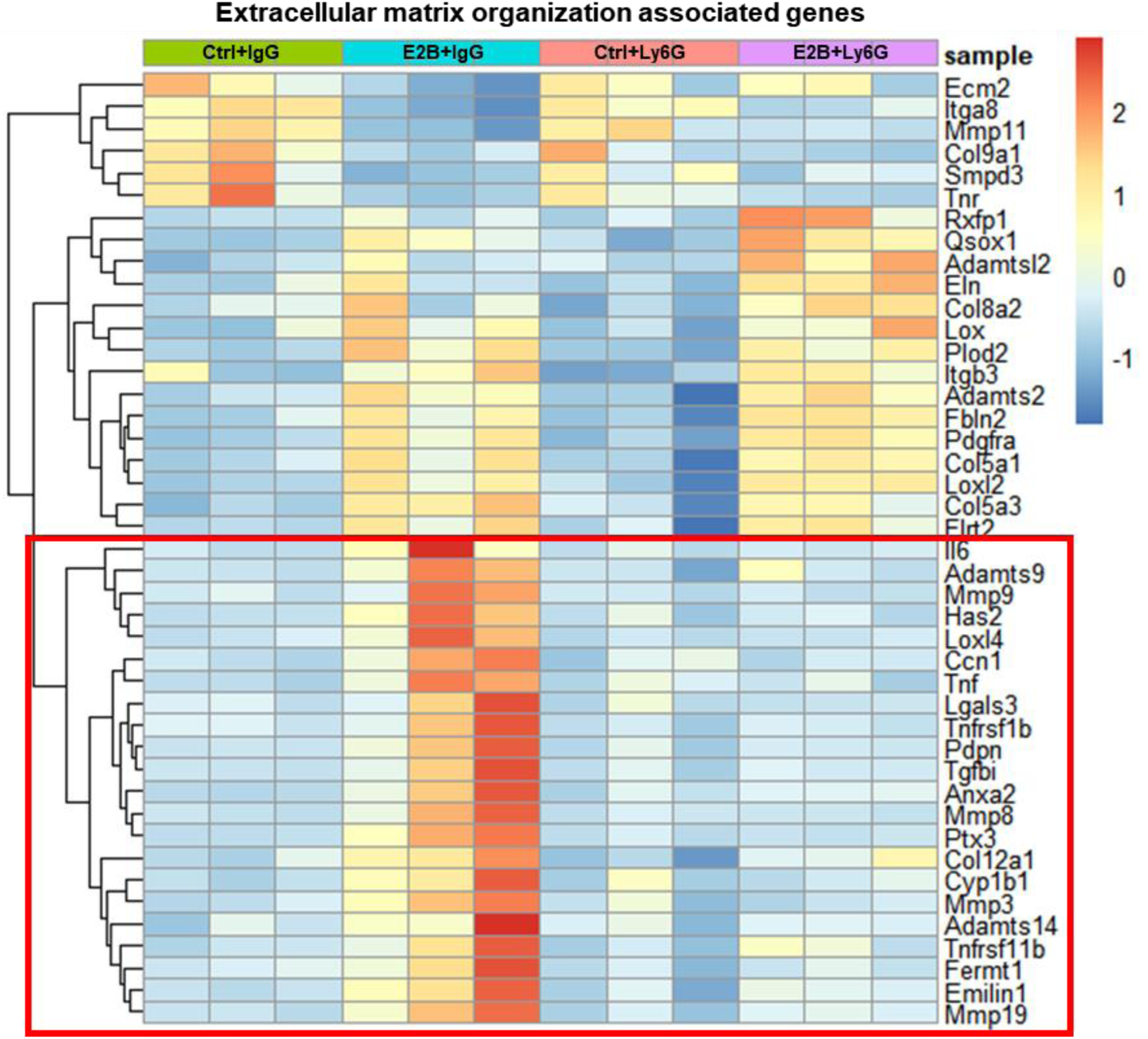
50% of estrogen-regulated ECM genes are abolished by neutrophil depletion. Mice at INV D1 was treated with anti-Ly6G antibody (Ly6G) or isotype control (IgG). 24h later, they were treated with vehicle control (Ctrl) or E2B for 24h. Heatmap representation of genes associated to extracellular matrix organization identified from the GO over-representation analysis (≥ 2 and ≤ −2-fold) of the DESeq2 analysed RNA-Seq data; Highlighted red box indicates part of the heatmap replotted and presented in Fig. 4C; CTRL+IgG n=3, CTRL+Ly6G n=3, E2B+IgG n=3, E2B+Ly6G n=3.

**Supplementary Figure 5.**
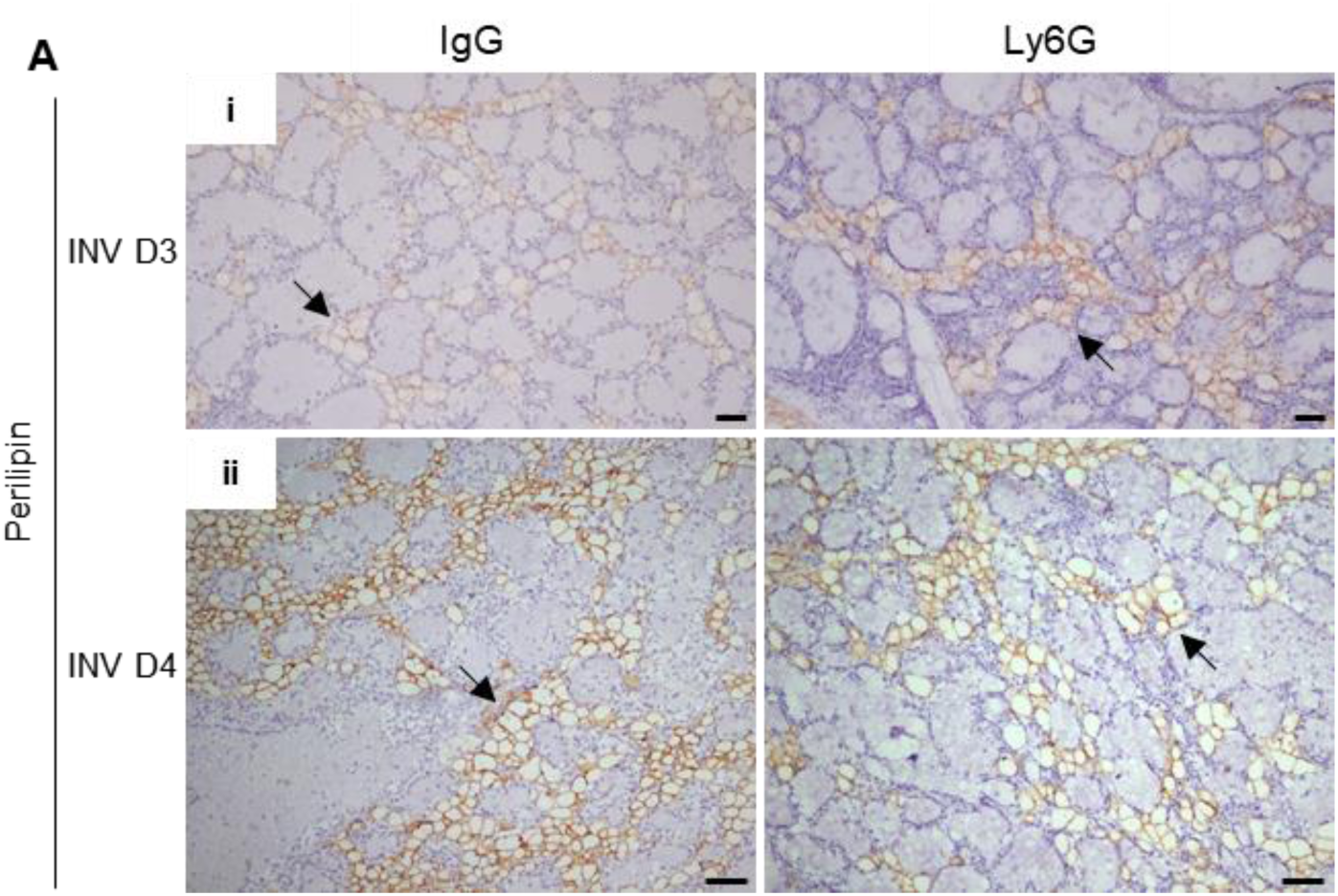
Neutrophil depletion transiently reduces E2B-induced adipocyte repopulation during mammary involution. Non-OVX mice were treated daily with either anti-Ly6G antibody (Ly6G) or isotype control (IgG) at 24h post-weaning (INV D1). Mammary tissue was collected for analysis at INV D3 and INV D4. A, (i) Perilipin IHC at INV D3; (ii) Perilipin IHC at INV D4; arrows indicate perilipin^+^ adipocytes. Scale bars: 50µm.

**Supplementary Figure 6.**
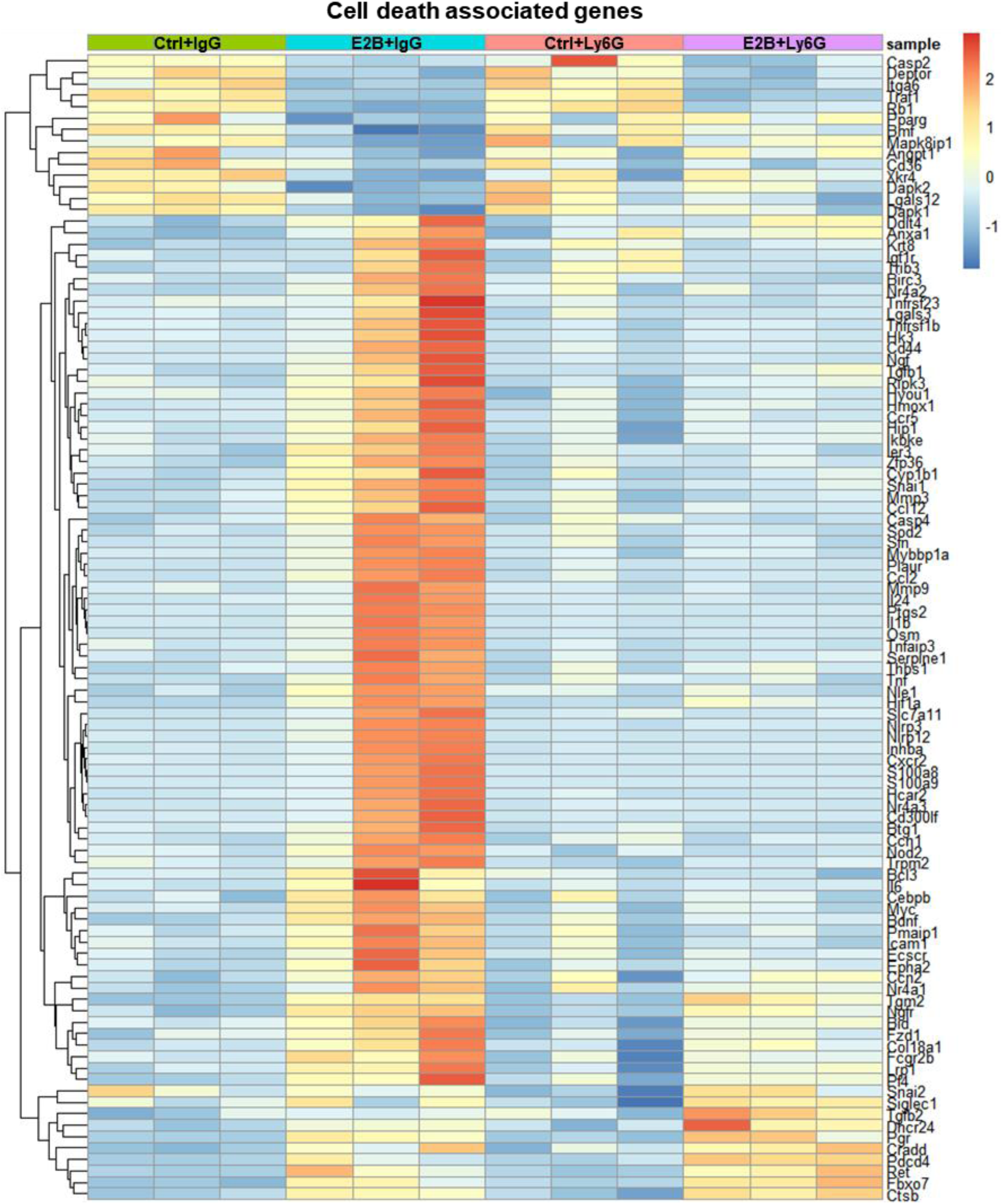
Heatmap representation of estrogen-regulated genes associated with cell death from the GO over-representation analysis. Mice at INV D1 was treated with anti-Ly6G antibody (Ly6G) or isotype control (IgG). 24h later, they were treated with vehicle control (Ctrl) or E2B for 24h. The genes plotted exhibit fold change of ≥ 1.5 and ≤ −1.5 with padj<0.05 from the DESeq2 analysis of the RNA-Seq data. Ctrl+IgG n=3, Ctrl+Ly6G n=3, E2B+IgG n=3, E2B+Ly6G n=3.

**Supplementary Figure 7.**
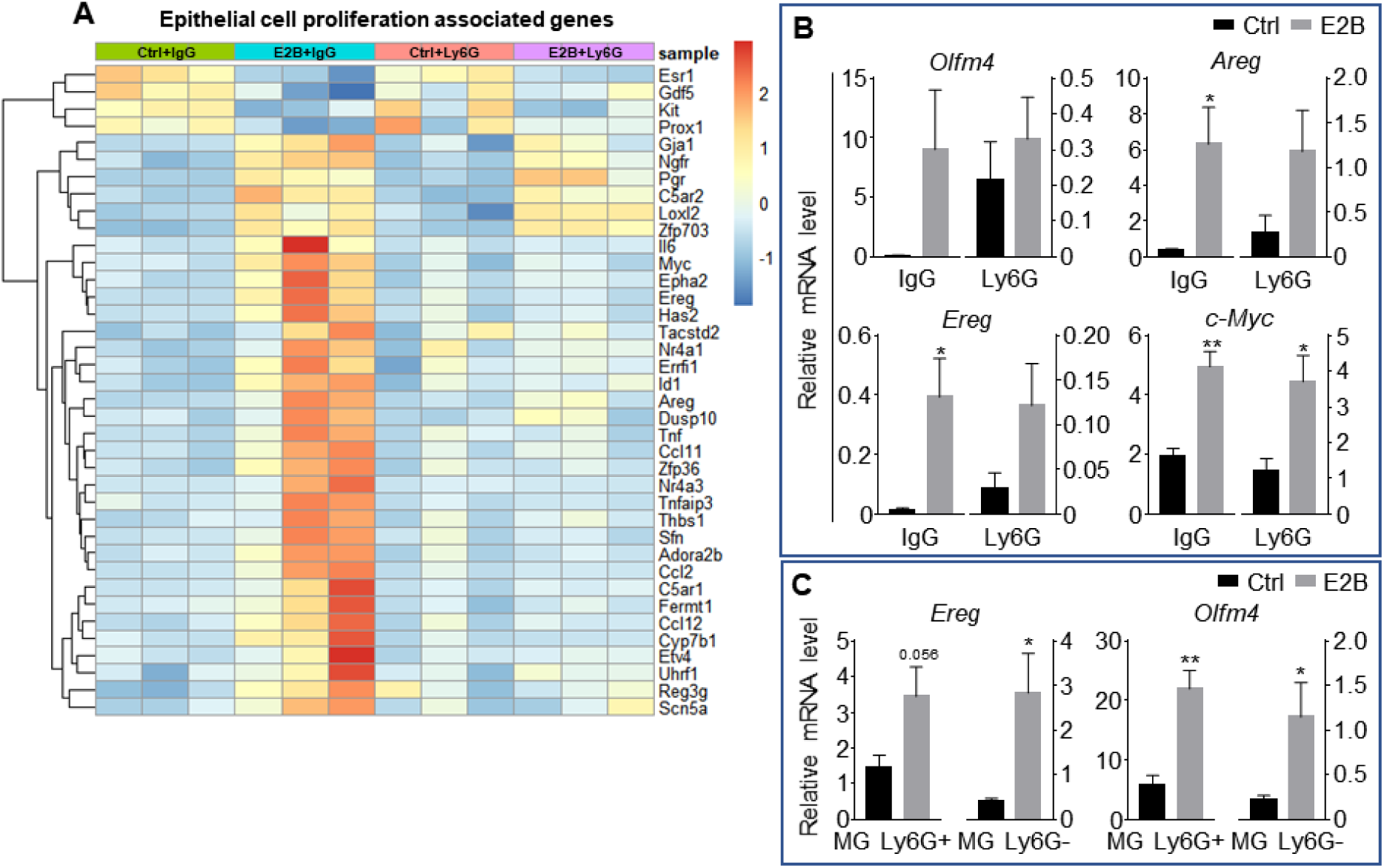
Estrogen-induced genes associated with cell proliferation are regulated in both mammary gland and neutrophil population. A-B, Mice at INV D1 was treated with anti-Ly6G antibody (Ly6G) or isotype control (IgG). 24h later, they were treated with vehicle control (Ctrl) or E2B for 24h; A, Heatmap representation of genes associated to epithelial cell proliferation identified from the GO over-representation analysis (≥ 2 and ≤ −2-fold) of the DESeq2 analysed RNA-Seq data; B, E2B induces proliferative gene expression independent of neutrophil presence; qPCR validation of estrogen-induced expression of *Areg*, *c-Myc*, and *Ereg* relative to *36b4* (Ctrl+IgG n=3, Ctrl+Ly6G n=3, E2B+IgG n=3, E2B+Ly6G n=3). C, *Ereg* and *Olfm4* were induced by estrogen in both mammary neutrophil and non-neutrophil population; Mice at INV D2 was treated with Ctrl or E2B for 24h. Gene expression of *Ereg* and *Olfm4* analysed in both Ly6G+ and Ly6G-population by qPCR analysis (Ctrl n=5, E2B n=5). Data represented as mean ± SEM.

**Supplementary Figure 8.**
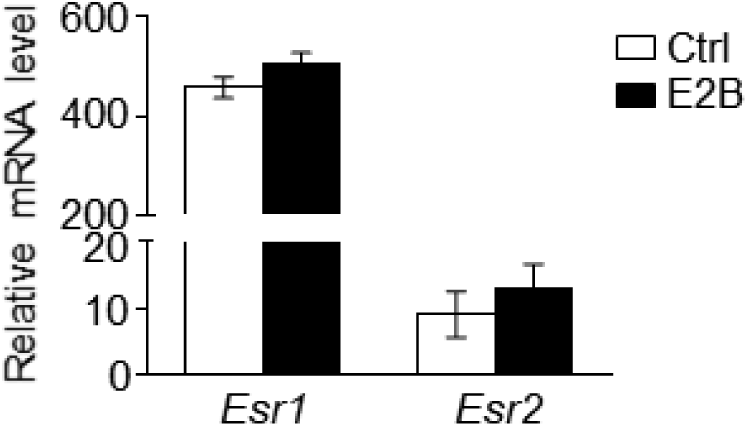
Expression of *Esr1* (ERα) is about 40 times higher than *Esr2* (ERβ) in mammary neutrophils during mammary involution. Mice at INV D2 was treated with either vehicle control (Ctrl) or E2B for 8h. Mammary neutrophils were isolated from tissue using Dynabeads® bound anti-Ly6G antibody. Gene expression of *Esr1* and *Esr2* relative to *36b4* were analysed in Ly6G+ population by qPCR analysis (Ctrl n=4, E2B n=4). Data represented as mean ± SEM.

**Supplementary Figure 9.**
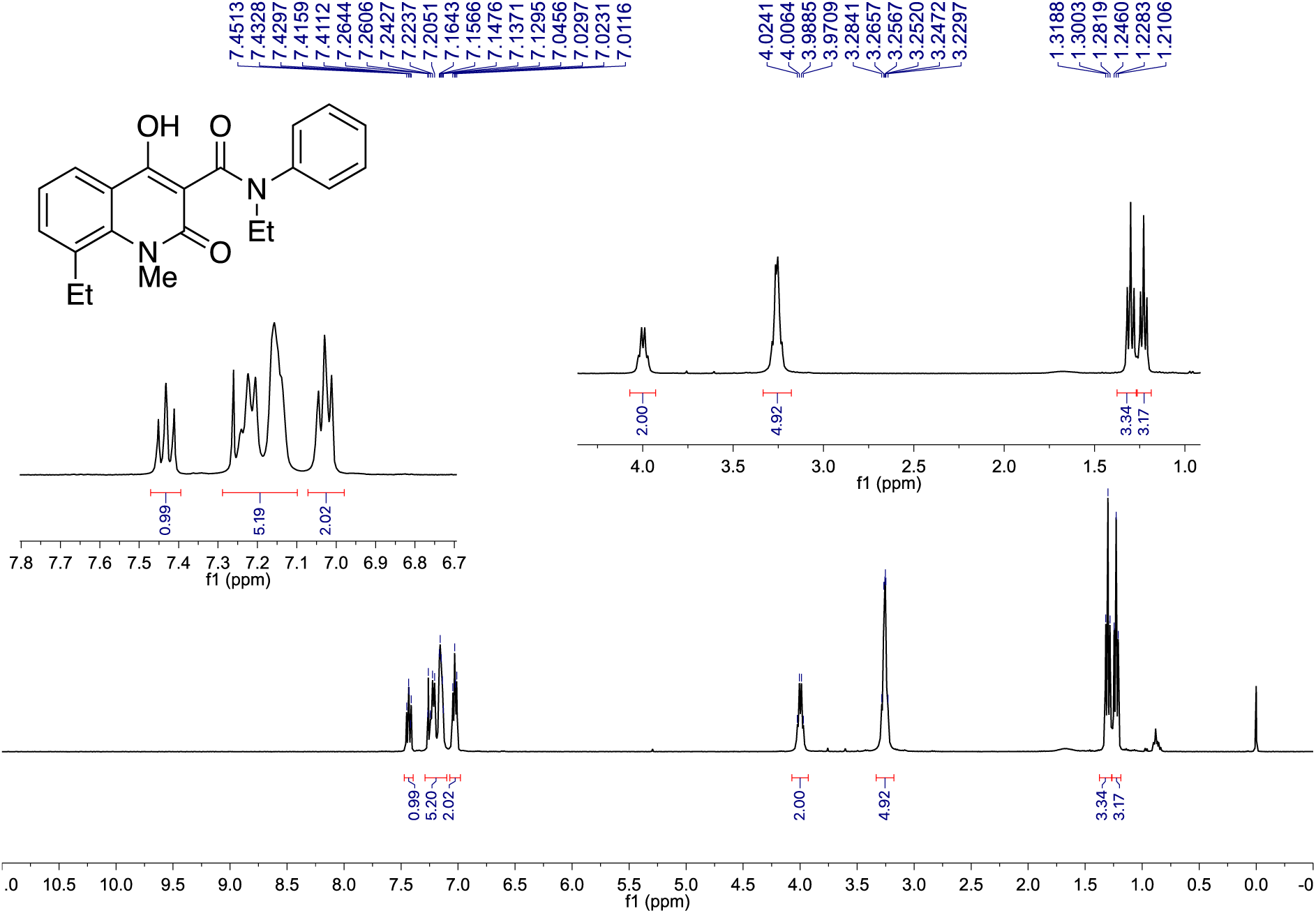
^1^H NMR Spectrum of the synthesized Paquinimod (PAQ): ^1^H NMR (CDCl3, 400 MHz) δ 7.43 (1H, dd, *J* = 8.6, 7.4 Hz), 7.29 – 7.10 (5H, m), 7.03 (2H, dd, *J* = 8.1, 5.5 Hz), 4.00 (2H, q, *J* = 7.1 Hz), 3.28 – 3.23 (3H+2H m), 1.30 (3H, t, *J* = 7.4 Hz), 1.23 (3H, t, *J* = 7.1 Hz). ^1^H NMR spectrum was recorded on a Bruker Avance 400 spectrometer.

**Supplementary Table 1.**
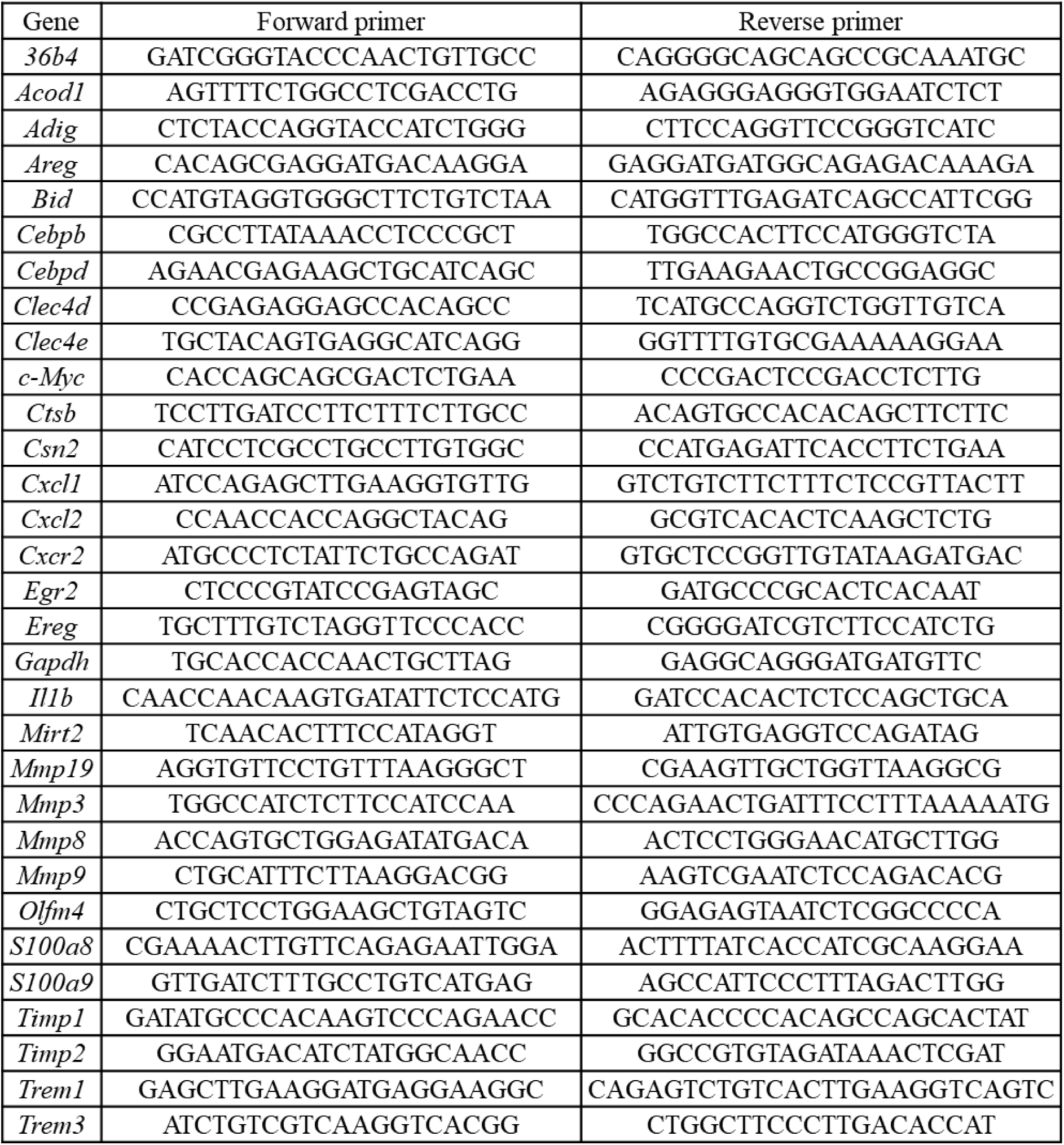
List of qPCR primers used.

**Supplementary Table 2.** List of antibodies used.

| Antibody | Catalog number | Dilution | Company |
| --- | --- | --- | --- |
| Mouse anti-STAT3 | 610189 | 1:1000 | BD Biosciences |
| Rabbit anti-p-STAT3 (Y705) | 9131 | 1:1000 | Cell Signaling |
| Goat anti-CTSD | Sc-6486 | 1:1000 | Santa Cruz Biotechnology |
| Goat anti-CTSB | Sc-6493 | 1:1000 | Santa Cruz Biotechnology |
| Rat anti-CTSL | MAB9521 | 1:1000 | RnD Systems |
| Rabbit anti-PARP | 9542 | 1:1000 | Cell Signaling |
| Mouse anti-p53 (Ab-6) | OP43 | 1:1000 | Calbiochem |
| Mouse anti-GAPDH | AM4300 | 1:10000 | Ambion |
| Rabbit anti-LAMP2 | ADI-905-723-100 | 1:1000 | Enzo Life Sciences |
| Mouse anti-β-tubulin | M2008 | 1:10000 | SABio |
| Anti-rabbit IgG, HRP-linked Antibody | 7074 | 1:2000 | Cell Signaling |
| Rabbit anti-Cleaved caspase 3 | 9661 | 1:1000 | Cell Signaling |
| Rabbit anti-Perilipin | 3470 | 1:100 | Cell Signaling |
| Anti-mouse IgG, HRP-linked Antibody | 7076 | 1:2000 | Cell Signaling |
| rabbit anti-goat IgG-HRP | sc-2768 | 1:5000 | Santa Cruz Biotechnology |

